# Multi-omics analysis reveals distinct non-reversion mechanisms of PARPi resistance in BRCA1- versus BRCA2-deficient mammary tumors

**DOI:** 10.1101/2022.09.07.506927

**Authors:** Jinhyuk Bhin, Mariana Paes Dias, Ewa Gogola, Frank Rolfs, Sander R. Piersma, Roebi de Bruijn, Julian R. de Ruiter, Bram van den Broek, Alexandra A. Duarte, Wendy Sol, Ingrid van der Heijden, Lara Bakker, Taina S. Kaiponen, Cor Lieftink, Ben Morris, Roderick L. Beijersbergen, Marieke van de Ven, Connie R. Jimenez, Lodewyk F. A. Wessels, Sven Rottenberg, Jos Jonkers

## Abstract

BRCA1 and BRCA2 both function in DNA double-strand break repair by homologous recombination (HR). Due to their HR-defect, BRCA1/2-deficient cancers are sensitive to poly(ADP-ribose) polymerase inhibitors (PARPi) but they eventually acquire resistance. Preclinical studies yielded several PARPi resistance mechanisms that do not involve BRCA1/2 reactivation, but their relevance in the clinic remains elusive. To investigate which BRCA1/2-independent mechanisms drive spontaneous resistance *in vivo*, we combined molecular profiling with functional analysis of the HR status of matched PARPi-naïve and PARPi-resistant mouse mammary tumors harboring large intragenic deletions that prevent functional restoration of BRCA1/2. We observed restoration of HR in 64% of PARPi-resistant BRCA1-deficient tumors but none in the PARPi-resistant BRCA2-deficient tumors. Moreover, we found that 53BP1 loss is the prevalent resistance mechanism in HR-proficient BRCA1-deficient tumors, whereas resistance in BRCA2-deficient tumors is mainly induced by the loss of PARG. Our combined multi-omics analysis catalogued additional genes and pathways potentially involved in modulating PARPi response.

## INTRODUCTION

The observation that many oncogenic events render cancer cells reliant on specific and druggable biological pathways is a premise of targeted therapies for personalized cancer treatment. Unfortunately, the selective pressure that initially kills cancer cells is also a driving force in selecting cells which acquired drug resistance. A better understanding of the recurrent molecular patterns of resistance in specific genetic contexts is therefore instrumental to improve clinical outcomes.

One example of cancer dependencies that can be exploited therapeutically is the defect in the repair of DNA double-strand breaks (DSBs) through homologous recombination (HR) due to BRCA1 or BRCA2 inactivation (Bryant et al., 2005; Farmer et al., 2005). Unlike the other DSB repair pathways, HR enables the accurate repair of DNA lesions as it uses the newly replicated sister chromatid as a template. Both BRCA1 and BRCA2 are essential in this process. While BRCA1 is required for the initiation of HR by promoting end-resection of the DSB, BRCA2 acts further downstream and, together with PALB2, stimulates the recruitment of RAD51 recombinase to the resected single-stranded DNA (Scully et al., 2019). The HR defect resulting from the loss of BRCA1/2 can be targeted through the inhibition of poly(ADP-ribose) polymerase (PARP) enzymes PARP1 and PARP2 (Bryant et al., 2005; Farmer et al., 2005). PARP1/2 have been implicated in several DNA damage response (DDR) pathways, including the repair of DNA single strand breaks (SSBs), DSBs and stabilization of replication forks (RFs) (Ray Chaudhuri and Nussenzweig, 2017). Catalytic inhibition as well as trapping of PARP1/2 on the DNA by PARP inhibitors (PARPi) leads to replication-coupled DSBs formation, which in turn requires HR for error-free repair (Murai et al., 2012). BRCA1/2-defective cells can only employ error-prone repair to resolve the DSBs caused by PARPi treatment, resulting in accumulation of chromosomal aberrations and cell death by mitotic catastrophe (Lupo and Trusolino, 2014). The success of this approach resulted in the clinical approval of four different PARPi for the treatment of several types of cancers with HR defects (Paes Dias et al., 2021).

Despite the clinical benefit, sustained antitumor responses to PARPi are often hampered by the emergence of resistance. Previous studies have delineated several mechanisms by which BRCA1/2-deficient tumors evade PARPi toxicity (Paes Dias et al., 2021). Independently of HR, PARPi resistance may be induced through (i) cellular extrusion of PARPi by upregulation of the drug efflux transporter P-gp (Rottenberg et al., 2007); (ii) partial restoration of catalytic PARP activity through loss of poly(ADP-ribose) glycohydrolase (PARG) (Gogola et al., 2018); (iii) PARP1 downregulation/inactivation as well as mutations that abolish PARP1 trapping (Murai et al., 2012; Pettitt et al., 2013, 2018); and (iv) restoration of RF stability (Cantor and Calvo, 2017; Murai et al., 2016, 2018). All these mechanisms result in PARPi resistance by limiting PARPi-induced DNA damage, rather than restoring the capacity of BRCA1/2-deficient cells to efficiently repair DSBs. In contrast, HR restoration as result of secondary (epi)genetic events that lead to reactivation of functional BRCA1/2 may fully cancel the initial susceptibility to PARPi. In addition, genetic screens and *in vivo* studies in preclinical models demonstrated that inactivation of the 53BP1-RIF1-shieldin DSB end-protection pathway, which inhibits HR and is antagonized by BRCA1 during S phase, partially restores HR and confers PARPi resistance in BRCA1-deficient cells (Boersma et al., 2015; Bouwman et al., 2010; Bunting et al., 2010; Chapman et al., 2013; Dev et al., 2018; Escribano-Díaz et al., 2013; Feng et al., 2015; Findlay et al., 2018; Gao et al., 2018; Ghezraoui et al., 2018; Gupta et al., 2018; Noordermeer et al., 2018; Tomida et al., 2018; Di Virgilio et al., 2013; Xu et al., 2015; Zimmermann et al., 2013).

While multiple mechanisms of acquired PARPi resistance have been reported in preclinical *in vitro* models, their clinical relevance remains unclear. To date, the best clinically documented mechanism of resistance is restoration of BRCA1/2 function by secondary (epi)genetic events (e.g., reversion mutations) (Ganesan, 2018). However, these results might be biased by the fact that PARPi were initially approved for second-line maintenance therapy following first-line treatment with platinum-based chemotherapies. Since (epi)genetic reactivation of BRCA1/2 function has been shown to be the main mechanism of platinum resistance in *BRCA1/2-mutated* tumors, it is plausible that some of these patients might have already developed BRCA-proficient, and therefore PARPi-resistant, tumor clones as a result of a first-line treatment (Barber et al., 2013; Domchek, 2017; Norquist et al., 2011; Weigelt et al., 2017). Moreover, reactivation of BRCA1/2 function is not found in all patients with refractory tumors (Ang et al., 2013; Weigelt et al., 2017), suggesting that BRCA1/2-independent PARPi resistance is relevant in the clinic.

The PARPi olaparib and niraparib have recently been approved as first-line maintenance therapies and clinical trials have started to test PARPi as single-agent neoadjuvant therapy (Litton et al., 2020). With more patients likely to receive PARPi earlier in the course of disease, it is important to understand which are the most frequent mechanisms of acquired PARPi resistance, and if these differ between *BRCA1-* and *BRCA2*-mutated tumors, in order to predict PARPi response and to develop strategies to overcome resistance. In the absence of available clinical data, we sought to answer these questions by combining functional analysis of HR status with molecular profiling of a unique collection of matched PARPi-naïve and PARPi-resistant mouse mammary tumors which harbor large intragenic deletions of *Brca1* or *Brca2* genes that cannot be spontaneously restored. Overall, our study shows that functional differences between BRCA1 and BRCA2 in the repair of DSBs also impact the resistance patterns in PARPi-treated tumors. While HR restoration accounted for the majority of BRCA1-deficient tumors, it did not occur in BRCA2-deficient tumors, suggesting that HR cannot be restored in *Brca2*-mutated tumors that cannot undergo BRCA2 reactivation. Moreover, among the previously reported resistance mechanisms, loss of 53BP1 and loss of PARG were the most dominant alterations in PARPi-resistant BRCA1-deficient and BRCA2-deficient tumors, respectively. Dysregulation of other known resistance factors was only sporadically observed, suggesting 53BP1 and PARG as potential biomarkers of acquired PARPi resistance. Additionally, our analysis yielded a list of potential genes and pathways involved in PARPi response and provides evidence that tumor-intrinsic alterations in pathways regulating the tumor microenvironment may influence PARPi efficacy.

## RESULTS

### HR restoration drives PARPi resistance in BRCA1-deficient tumors

To study the contribution of BRCA1/2-independent PARPi resistance mechanisms in BRCA1/2-deficient tumors, we used two genetically engineered mouse models (GEMMs) of BRCA1 - associated breast cancer, *K14cre;Brca1^F/F^;Trp53^F/F^* (KB1P) and *K14cre;Brca1^F/F^; Trp53^F/F^;Mdr1a/b^-/-^* (KB1PM) as well as a GEMM of BRCA2-associated breast cancer, *K14cre;Brca2^F/F^; Trp53^F/F^* (KB2P) (Gogola et al., 2018; Jaspers et al., 2013; Jonkers et al., 2001; Liu et al., 2007; Rottenberg et al., 2007) (**Figure 1A**). In these models, long-term treatment of mammary tumors with PARPi leads to spontaneous induction of resistance, which is preserved upon tumor passaging (Jaspers et al., 2013; Ray Chaudhuri et al., 2016). Importantly, the tumors arising in these models harbor large intragenic deletions in the *Brca1* or *Brca2* genes (Jonkers et al., 2001; Liu et al., 2007) and thus resistance to PARPi cannot be acquired via reactivation of BRCA1/2 function. Moreover, we eliminated the possibility of P-glycoprotein (Pgp)-mediated resistance to the PARPi olaparib by either the genetic inactivation of Pgp *(Mdr1a/b)* in the KB1PM model, or by treating Pgp-proficient KB1P and KB2P tumors with the PARPi AZD2461, which is a poor substrate for this transporter (Jaspers et al., 2013; Oplustil O’Connor et al., 2016) (**Figure 1A**).

**Figure 1.**
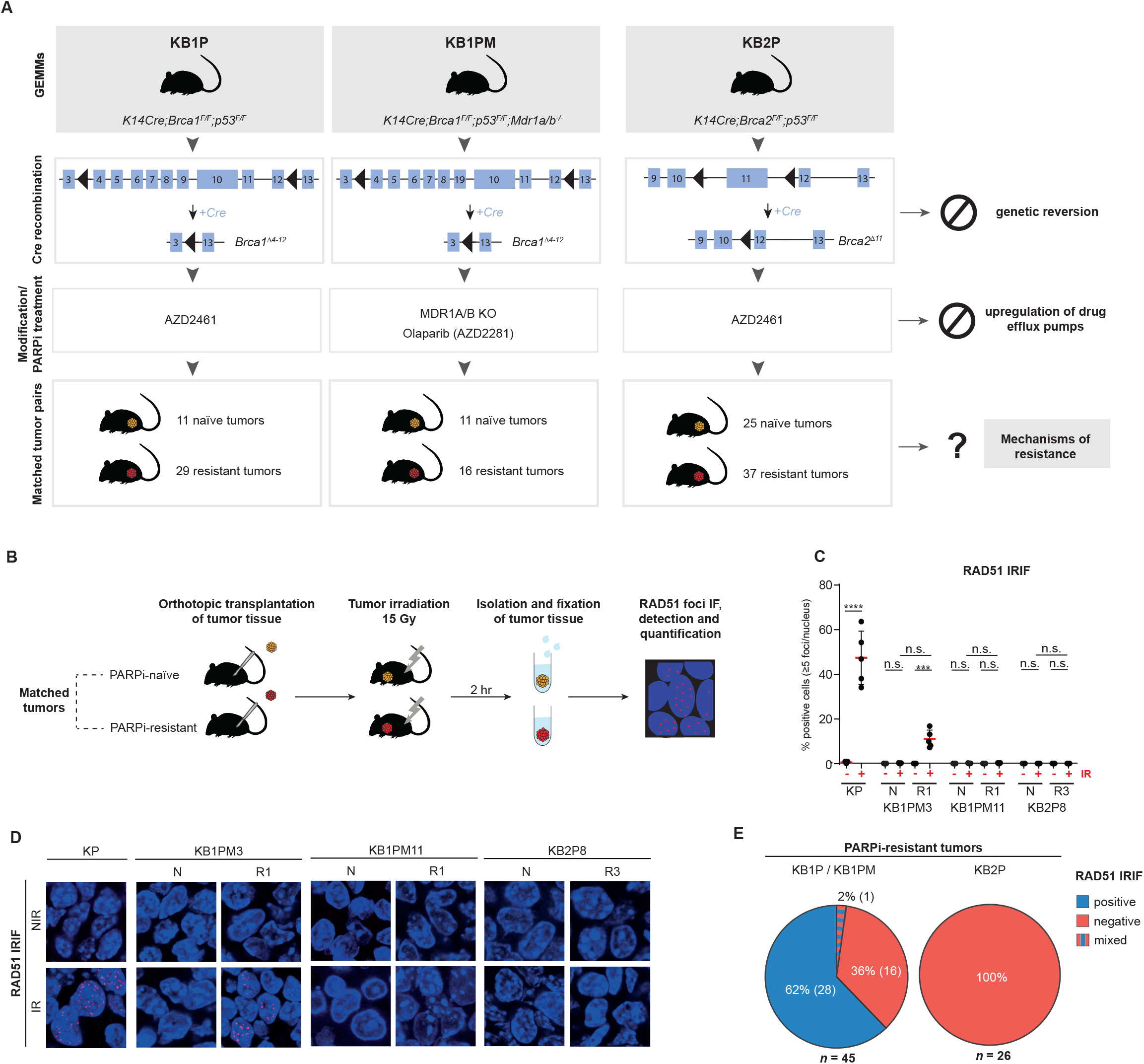
HR restoration drives PARPi resistance in BRCA1-deficient tumors. **(A)** Outline of the generation of matched PARPi-naive and PARPi-resistant KB1P(M) and KB2P tumors and of the experimental approach. **(B)** Schematic representation of the RAD51 IRIF formation assay. Cryopreserved PARPi-naïve and PARPi-resistant tumors were orthotopically transplanted into syngeneic recipient mice and, upon outgrowth to 500 mm3, DNA damage was inflicted by locally applied ionizing radiation (IR) at a dose of 15Gy. 2hr post-irradiation, tumors were isolated and fixed tissues were used for RAD51 immunofluorescence imaging. **(C-D)** Quantification (C) and representative images (D) of the RAD51 IRIFs for the different matched KB1P(M) and KB2P tumor pairs and control KP tumors; IR – irradiated, NIR – non irradiated; scale bar, 100 μm; data in (C) represented as percentage of positive cells (≥5 foci/nucleus) per imaged area (single data point, typically 100-200 cells/area); ****p<0.0001, ***p<0.001, ** p<0.01 (two-tailed Mann-Whitney U test). **(E)** Pie charts showing the outcome of the RAD51 IRIF assay in PARPi-resistant KB1P(M) and KB2P tumor cohorts; percentages and numbers of individual tumors analyzed are indicated; n – total number of individual tumors analyzed from the indicated models. p=0.0001 (two-tailed Fisher’s exact test).

HR deficiency is the basis for sensitivity to PARPi and thus we hypothesized that HR restoration is the most likely way for tumors to acquire resistance to PARPi. We therefore assessed the HR status of matched PARPi-naïve and PARPi-resistant tumors by measuring their capacity to form ionizing radiation-induced RAD51 foci (RAD51-IRIF) (Godin et al., 2016; Xu et al., 2015) (**Figure 1B**). As a positive control for this assay, we used a HR-proficient mammary tumor derived from the *K14cre;Trp53^F/F^* (KP) model and observed the highest accumulation of RAD51 foci 2 hours after induction of DNA damage (**Figures S1A-S1D**). Of note, all tumors exhibited high growth rates prior to irradiation, suggesting that differences in cell cycle distribution between the samples are negligible (**Figures S1E and S1F**). As expected, we did not detect any RAD51-IRIF formation in any of the PARPi-naïve tumors (**Figures 1C and 1D**), confirming that the *Brca1/2* deletions induced in our models completely abolish HR-mediated repair. Consistent with this, whole-exome sequencing of DNA from PARPi-resistant tumors confirmed the complete Cre-mediated deletion of the floxed *Brca1/2* exons, thus no emerging resistance could be attributed to the selection of clones that retained wild-type BRCA proteins (**Figures S1G and S1H**). Analysis of the 45 PARPi-resistant BRCA1-deficient (BRCA1-KO) tumors revealed that 64% (29/45) of the tumors had restored the capacity to form RAD51 foci, including one tumor with a mixed pattern (RAD51-IRIF positive and negative areas) (**Figure 1E**). These results indicate that HR recovery is the predominant mechanism of PARPi resistance in the KB1P(M) models, albeit not the only one. In contrast, none of the 26 PARPi-resistant BRCA2-deficient tumors exhibited RAD51-IRIF (**Figure 1E**) (Gogola et al., 2018). Given that PARPi treatment is a potent trigger of HR restoration in the KB1P(M) models, lack of RAD51-IRIF in the BRCA2-deficient cohort strongly indicates that BRCA2 is indispensable for HR repair.

### Alterations in previously reported PARPi resistance factors

To understand how prolonged PARPi treatment reshapes BRCA1/2-deficient tumors, we performed whole-exome sequencing (WE-seq), low-coverage whole-genome sequencing (LCWG-seq), and RNA sequencing (RNA-seq) on the collection of matched PARPi-naïve and PARPi-resistant KB1P(M) and KB2P tumors and identified alterations specific to each resistant tumor compared to the matched naïve tumor.

We first interrogated if we could find genetic and transcriptional alterations in factors previously associated with PARPi resistance (**Figure 2A**). To this end, we selected 25 genes reported to drive PARPi resistance due to BRCA-independent HR restoration, restoration of fork stability or modulation of PARP signaling/trapping (Paes Dias et al., 2021) (**Table S1**). We examined alterations in these genes in the different PARPi-resistant tumor groups with informed HR status, i.e., (1) RAD51-positive KB1P(M), (2) RAD51-negative KB1P(M) and (3) KB2P tumors. Of the 25 genes, 23 have been reported to drive PARPi resistance upon loss of function, whereas 2 genes drive resistance as a result of gain of function. Globally, we found alterations in all known factors analyzed, which occurred at different frequencies in the different PARPi-resistant tumor groups. Nearly eighty percent (55/71) of all resistant tumors harbored deleterious mutations, copy number variations, and/or gene expression changes in at least one known factor. *Shld2, Parg, Rif1, Trp53bp1, Rev7, Ezh2, Mre11a, Mll3* and *Mll4* were amongst the most frequently altered genes (≥10% of all tumors) (**Figure 2A**).

**Figure 2.**
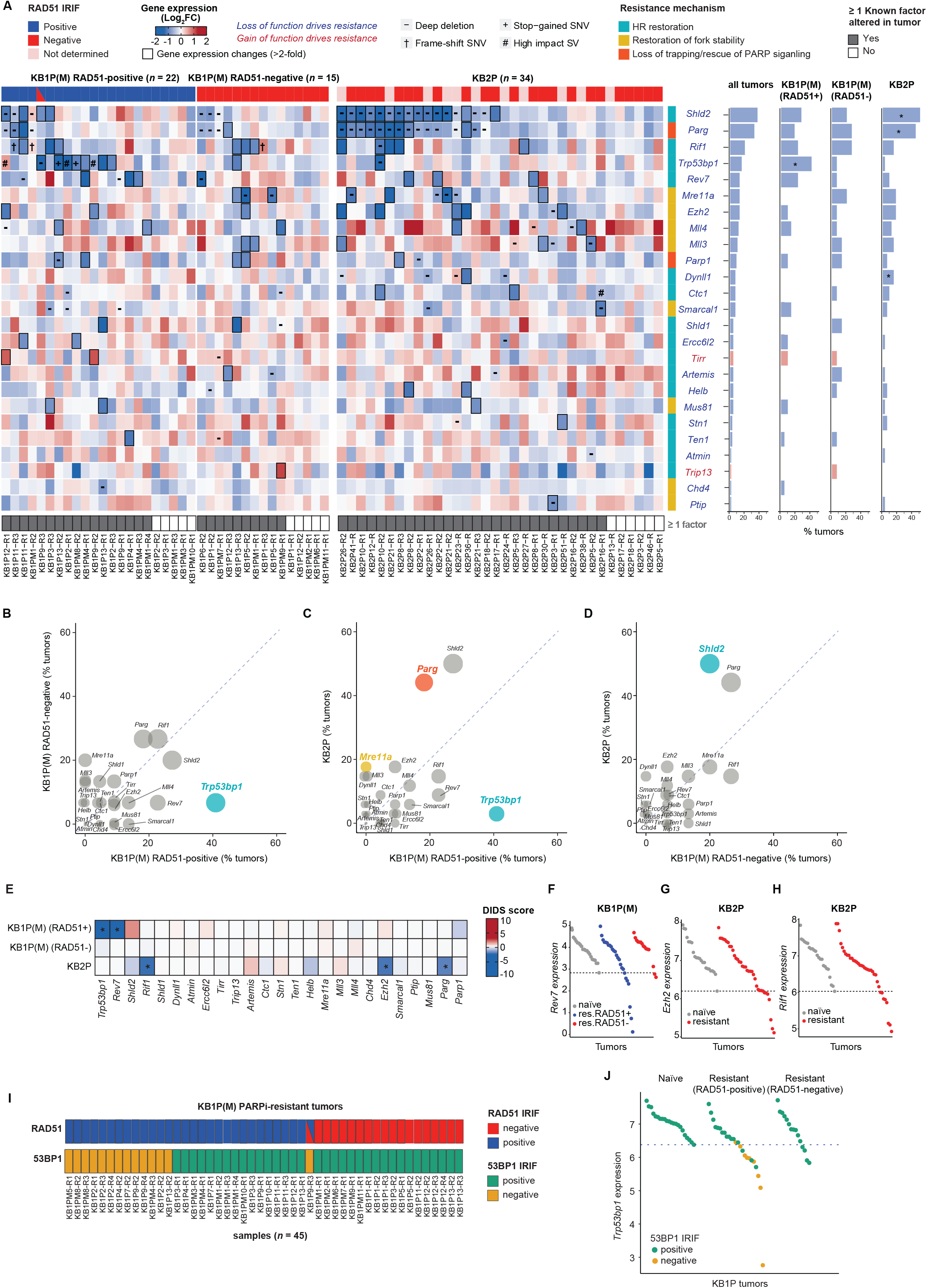
Alterations in previously reported PARPi resistance factors. **(A)** Heatmap (left) of gene expression changes between matched resistant vs naïve KB1P(M) and KB2P tumors for the previously reported PARPi resistance genes (Paes Dias et al., 2021). Genes for which loss or gain of function have been reported to drive resistance are indicated in blue or red, respectively. In the heatmap, genomic are marked by different symbols (-: copy number deletion, +: stop-gained SNV, †: frame-shift SNV, #: loss of function SV). The resistant tumors with transcriptional alterations (lower or higher than 2-fold compared to matched naïve tumors for each loss or gain of function drivers) are marked by thicker borderlines. The top panel of the heatmap indicates RAD51 IRIF status which is a proxy for HR status (positive - blue, negative - red, tumors with both positive and negative area - blue and red, tumors for which RAD51 IRIF was not determined, even though expected to be negative - pink). The bottom panel of the heatmap indicates the tumors with alterations in at least one known gene. Resistance mechanisms associated with each gene are categorized by different color bars (light green - HR restoration, purple – restoration of fork stability, light blue – PARP signaling). Frequencies for dysregulation of each gene (by either genomic or transcriptional alterations) are shown in the bar plots next to the heatmap (right). The genes preferentially altered in specific tumor types were assessed by the Fisher’s exact test (asterisk, P<0.05). **(B-D)** Scatter plots comparing the alteration frequency of each PARPi resistance factor in the different resistant tumor types. The size of the circle is proportional to the sum of the alteration frequency of the two resistant tumor types compared and circles are colored if statistically significant (Fisher’s exact test, P<0.05). The color of the circle indicates the resistance mechanism associated with each resistance factor as mentioned in Figure 2A. **(E)** DIDS outlier scores computed from gene expression data for known resistance factors. Red (positive score) and blue (negative score) indicate upregulation and downregulation of each factor in subset of resistant tumors compared to naïve tumors. The genes with significant DIDS scores are marked with asterisk (Permutation-based exact test, P<0.05). **(F-H)** Dot plots of *Rev7* gene expression in KB1P(M) tumors (F) and *Ezh2* (G) and *Rif1* (H) gene expressions in KB2P tumors where significant DIDS scores were detected. **(I)** RAD51 and 53BP1 IRIF status in KB1P(M) PARPi-resistant tumors measured by in situ IRIF assay. **(J)** Dot plots of *Trp53bp1* gene expression in KB1P(M) tumors with 53BP1 IRIF status.

We next asked whether certain genes were preferentially altered within the three different PARPi-resistant tumor groups. We found that *Trp53bp1*, one of the best-studied PARPi-resistance factors involved in HR restoration via loss of DSB end-protection, was specifically altered in RAD51-positive KB1P(M) tumors (41%) as compared to RAD51-negative KB1P(M) (7%) and KB2P (3%) tumors (**Figures 2A–2D**). In contrast, *Shld2* and *Parg* were preferentially altered by copy number loss in KB2P tumors compared to KB1P(M) tumors (**Figure 2A and 2C**). Previously, we reported that loss of PARG causes PARPi resistance independently of BRCA1/2 by restoring PAR formation and restoring the recruitment of DNA repair factors downstream of PARP1 (Gogola et al., 2018). While we expected *Parg* to be equally altered in both KB1P(M) and KB2P tumors, we found that *Parg* was more frequently lost in KB2P tumors (44%) (**Figure 2A and 2C**) than in RAD51-positive (18%) and RAD51-negative KB1P(M) tumors (27%) (**Figure 2B**). The genomic location of *Shld2* is proximal to *Parg* (chr14qB) and thus copy number loss of *Parg* is often accompanied by concomitant loss of *Shld2* (**Figure 2A**). Moreover, depletion of SHLD2 has been reported to drive PARPi resistance via HR restoration in BRCA1-deficient cells but not in BRCA2-deficient cells. Hence, loss of *Shld2* in KB2P tumors is most likely a consequence of *Parg* copy number loss rather than driving resistance to PARPi. In support of this, CRISPR/Cas9-mediated inactivation of SHLD2 did not drive resistance in cultured KB2P tumor cells (**Figures S2A and S2B**). MRE11 downregulation or copy loss was only found in KB2P and RAD51-negative KB1P(M) tumors, suggesting MRE11 is specifically lost in RAD51-negative PARPi-resistant tumors (**Figures 2A–2D**). This is in line with its key role in HR and suggests that loss of MRE11 may induce PARPi resistance by promoting RF protection.

We also applied Detection of Imbalanced Differential Signal (DIDS) analysis specifically designed for the detection of subgroup markers in heterogeneous populations (de Ronde et al., 2013). This tool is particularly useful for identifying drug resistance factors in a tumor group with multiple resistance mechanisms by detecting genes with outlying expression in the subset of resistant tumors compared to all naïve tumors (Gogola et al., 2018; Kas et al., 2018). DIDS analysis additionally identified *Rev7* to be significantly downregulated in RAD51-positive PARPi-resistant KB1P(M) tumors, which is consistent with its role in the 53BP1-RIF1-shieldin pathway involved in DSB end-protection, and with the observation that REV7 loss promotes HR and PARPi resistance in BRCA1-deficient cells (Boersma et al., 2015; Ghezraoui et al., 2018; Gupta et al., 2018; Noordermeer et al., 2018; Xu et al., 2015) (**Figures 2E and 2F**). *Ezh2*, which was previously shown to promote fork stability and PARPi resistance when depleted, was also identified by DIDS analysis to be downregulated in PARPi-resistant KB2P tumors, in accordance with the previous findings that loss of EZH2 impairs response to PARPi in BRCA2-deficient cells but not in BRCA1-deficient cells (Rondinelli et al., 2017) (**Figures 2E and 2G**). Surprisingly, *Rif1* was significantly downregulated in PARPi-resistant KB2P tumors, suggesting RIF1 might have 53BP1-RIF1-shieldin-independent functions that could drive resistance in BRCA2-deficient tumors (**Figures 2E and 2H**).

To confirm that PARPi resistance mediated by 53BP1 loss is enriched in RAD51-positive PARPi-resistant tumors, we evaluated the functional impairment of 53BP1 in KB1P(M) and KB2P tumors by analyzing 53BP1-IRIF formation (**Figures S2C and S2D**). We found that loss of 53BP1-IRIF was only detected in RAD51-positive or mixed PARPi-resistant KB1P(M) tumors, whereas PARPi-resistant KB2P tumors as well as PARPi-naïve KB1P(M) or KB2P tumors did not show loss of 53BP1-IRIF (**Figures 2I and S2E**). Consistently, tumors with loss of 53BP1-IRIF showed lower *Trp53bp1* expression levels than other tumors, suggesting good concordance between the omics data and the functional assay (**Figure 2J**). Altogether, our analysis revealed multiple known factors significantly altered in RAD51-positive KB1P(M) and in KB2P tumors; however, none were found to specifically explain PARPi resistance in RAD51-negative PARPi-resistant KB1P(M) tumors.

### HR-deficient PARPi-resistant BRCA1-KO tumors show increased immune cell infiltration

We then systematically characterized the three resistant tumor groups beyond the known resistance factors using our genomic and transcriptomics data. To identify recurrent focal genomic alterations between resistant and matched naïve tumors, we performed copy number analysis using RUBIC (van Dyk et al., 2016). The majority of the significantly recurrent alterations identified in PARPi-resistant tumors were focal deletions (**Figures S3A-S3C**), including loss of the regions encoding *Rev7* in RAD51-positive KB1P(M) tumors and *Parg* in KB2P tumors, respectively (**Figures S3A and S3C**). In PARPi-resistant RAD51-negative KB1P(M) tumors, we could not identify recurrent focal events, with the exception of one amplified locus on chromosome 8 (**Figure S3B**). Several genes encoded within the recurrently amplified/deleted loci in PARPi-resistant KB1P(M) and KB2P tumors are associated with DDR pathways (**Figure S3D**). Gene set analysis (GSA) demonstrated that genes involved in cell cycle/proliferation were depleted in PARPi-resistant RAD51-positive KB1P(M) tumors, whereas genes involved in positive regulation of immune cell activation were amplified in PARPi-resistant RAD51-negative KB1P(M) tumors (**Figures S3E-S3G**).

Next, we transcriptionally characterized the different PARPi-resistant tumor groups. Differential gene expression analysis by limma (Ritchie et al., 2015) between resistant and naïve tumors identified 26, 349 and 1058 genes in PARPi-resistant RAD51-positive KB1P(M), RAD51-negative KB1P(M) and KB2P tumors, respectively, including downregulation of *Trp53bp1, Parg* and *Shld2* (**Figures 2A and S3H-S3J**). No known resistance-associated factors were found to be differentially expressed in RAD51-negative KB1P(M) tumors (**Figure S3I**). GSA of the differentially expressed genes (DEGs) in each group of resistant tumors identified distinct sets of pathways. Interestingly, PARPi-resistant RAD51-negative KB1P(M) tumors showed upregulation of immune-related pathways, including antigen processing and presentation, T-cell receptor signaling and phagosome (**Figure 3A**), which might be associated with the enrichment of immune cell regulators with focal amplification in these tumors (**Figure S3F**). Co-functionality network analysis of upregulated immune-associated genes using the GenetICA-Network framework (Bhattacharya et al., 2020) revealed several immune cell modules, such as T cells (e.g., CD3D, CD3G, CD247), B cells (e.g., CD22, CD72, CD79A) and antigen presentation (e.g., CD74, CIITA, H2-OB, H2-AA, H2-AB1, H2-EB1, H2DMA, H2-DMB1, H2-DMB2), suggesting an increase in immune cell infiltration (**Figure 3B**). We validated these findings by carrying out immunohistochemical (IHC) analysis of markers of leucocytes (CD45), T cells (CD3), B cells (B220) and macrophages(F4/80), revealing increased expression of all these markers in PARPi-resistant KB1P(M) tumors when compared to naïve tumors, with RAD51-negative KB1P(M) tumors displaying a stronger increase compared to RAD51-positive KB1P(M) tumors (**Figure 3C**). Therefore, our findings suggest that RAD51-negative PARPi-resistant KB1P(M) tumors have higher immune infiltration compared to PARPi-naïve and RAD51-positive PARPi-resistant KB1P(M) tumors. Altogether, these data demonstrate that PARPi-resistant tumors display distinct genomic and transcriptomic features depending on BRCA1/2 loss and HR status.

**Figure 3.**
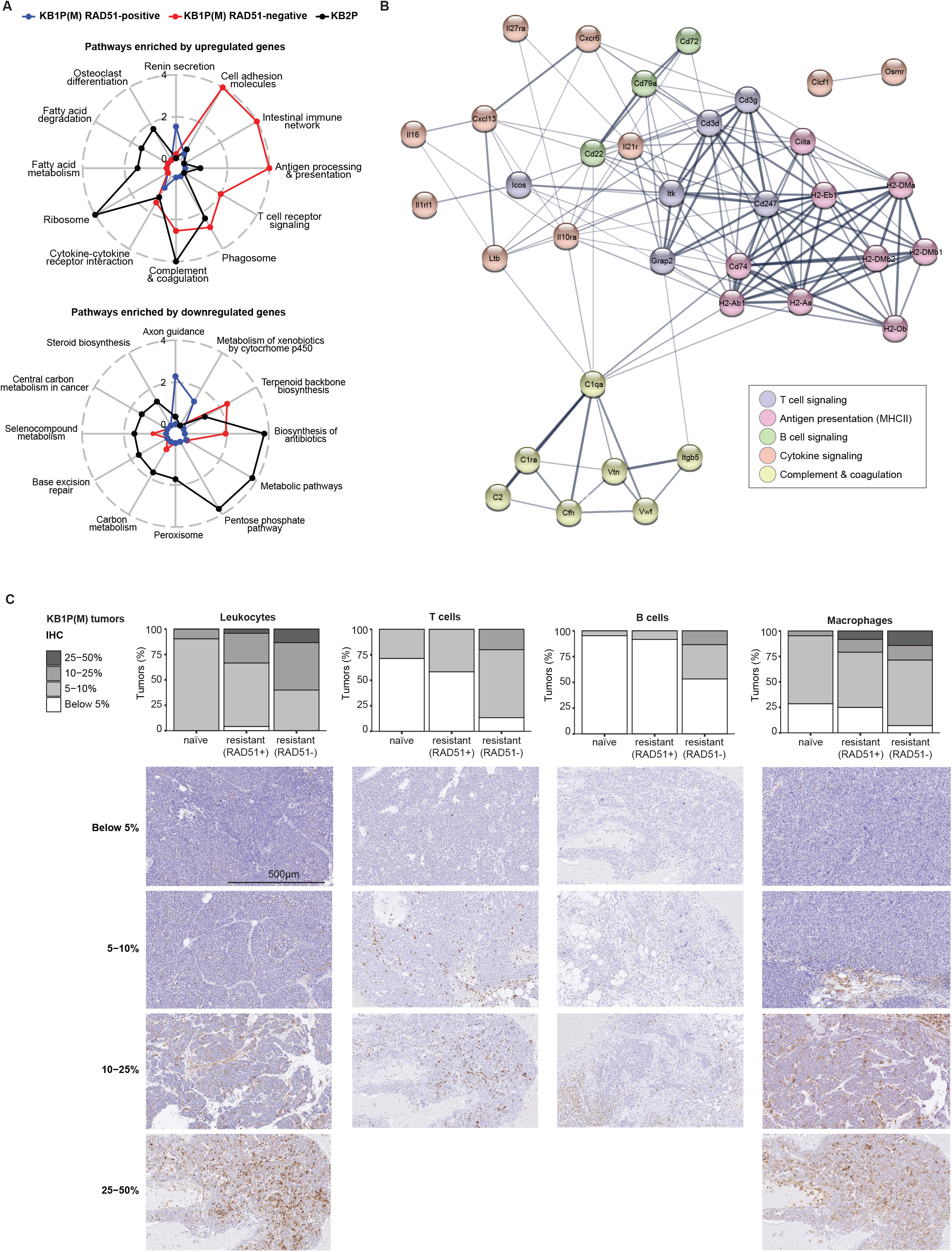
HR-deficient PARPi-resistant BRCA1-KO tumors show increased immune cell infiltration. **(A)** Radar chart showing pathways enriched by upregulated and downregulated genes in KB1P(M) and KB2P resistant tumors compared to naïve tumors based on gene expression data. **(B)** Co-functionality network constructed by STRING (Szklarczyk et al., 2021) for the immune-related genes that are significantly upregulated in RAD51-negative KB1P(M) resistant tumors compared to naive tumors. The genes in the network were annotated by KEGG (Szklarczyk et al., 2021) and colored depending on the annotated pathways. **(C)** Quantification and representative images of IHC analysis of markers for different immune cells including leukocytes (CD45), T-cells (CD3), B-cells (B220), and macrophages (F4/80) in KB1P(M) tumors.

### Multi-omics analysis identifies potential PARPi-resistance factors/pathways

To catalogue potential PARPi resistance factors, we identified genes displaying resistance-specific genomic (SNVs, INDELs, SVs, focal amplifications/deletions) and transcriptional (DEG sets) alterations in each group of resistant tumors by integrating WE-seq, LCWG-seq and RNA-seq data (**Figure 4A**). Of note, we extended DEG sets to capture the genes with both homogeneous (by limma) and heterogenous behavior (by DIDS) between naïve and resistant tumors. Overall, we observed limited overlap between genomic and transcriptional alterations across all tumor groups, including several genes involved in DDR (**Figure 4B; Table S2**). Moreover, the overlap between genomic and transcriptional alterations was mostly derived from copy number alterations, with the exception of *Trp53bp1*, in which we found truncating SNVs and deleterious SVs leading to nonsense-mediated decay. The limited overlap between genomic and transcriptional alterations indicates that the genomic alterations do not always lead to gene expression changes. Nonetheless, several known resistance factors including *Trp53bp1, Rev7*, and *Parg* displayed both genomic and transcriptional alterations. We next explored biological pathways represented by the identified genes with resistance-specific genomic and transcriptional alterations (**Figures 4C and 4D; Table S3**). As described before, we found transcriptional upregulation of immune-associated pathways specifically in RAD51-negative KB1P(M) tumors, including upregulation of antigen processing and presentation, B and T cell receptor signaling, as well as immune checkpoint pathways (**Figures 4C and 4D**). Interestingly, pathways associated with cell adhesion molecules and ECM-receptor interaction were transcriptionally downregulated in RAD51-positive KB1P(M) tumors, but upregulated in RAD51-negative KB1P(M) tumors (**Figures 4C and 4D**). At the genomic level, loss-of-function alterations (mainly copy number loss or deleterious SNVs/SVs) were observed in pathways associated with signaling processes such calcium and cAMP signaling pathways in RAD51-positive KB1P(M) tumors, and in metabolic pathways such as inositol phosphate and carbon metabolism in KB2P tumors (**Figures 4C and 4D**). In line with the limited overlap between genomic and transcriptional alterations (**Figure 4B**), pathways enriched by genes with genomic alterations showed very limited overlap with those enriched in transcriptional alterations, with the exception of the phagosome pathway (**Figure 4C**). Nonetheless, we found 46 genes that carried either genomic or transcriptional alterations in all three resistant tumor groups, including 8 genes *(Parp3, Gstm1, Il18, Padi4, Dnmt3b, Psrc1, Rif1* and *Ankrd26*) involved in DDR (**Figures 4E and 4F**). Taken together, integration of genomics and transcriptomics data from PARPi-resistant versus PARPi-naïve tumors allowed us to catalogue genes and pathways potentially involved in modulating PARPi response.

**Figure 4.**
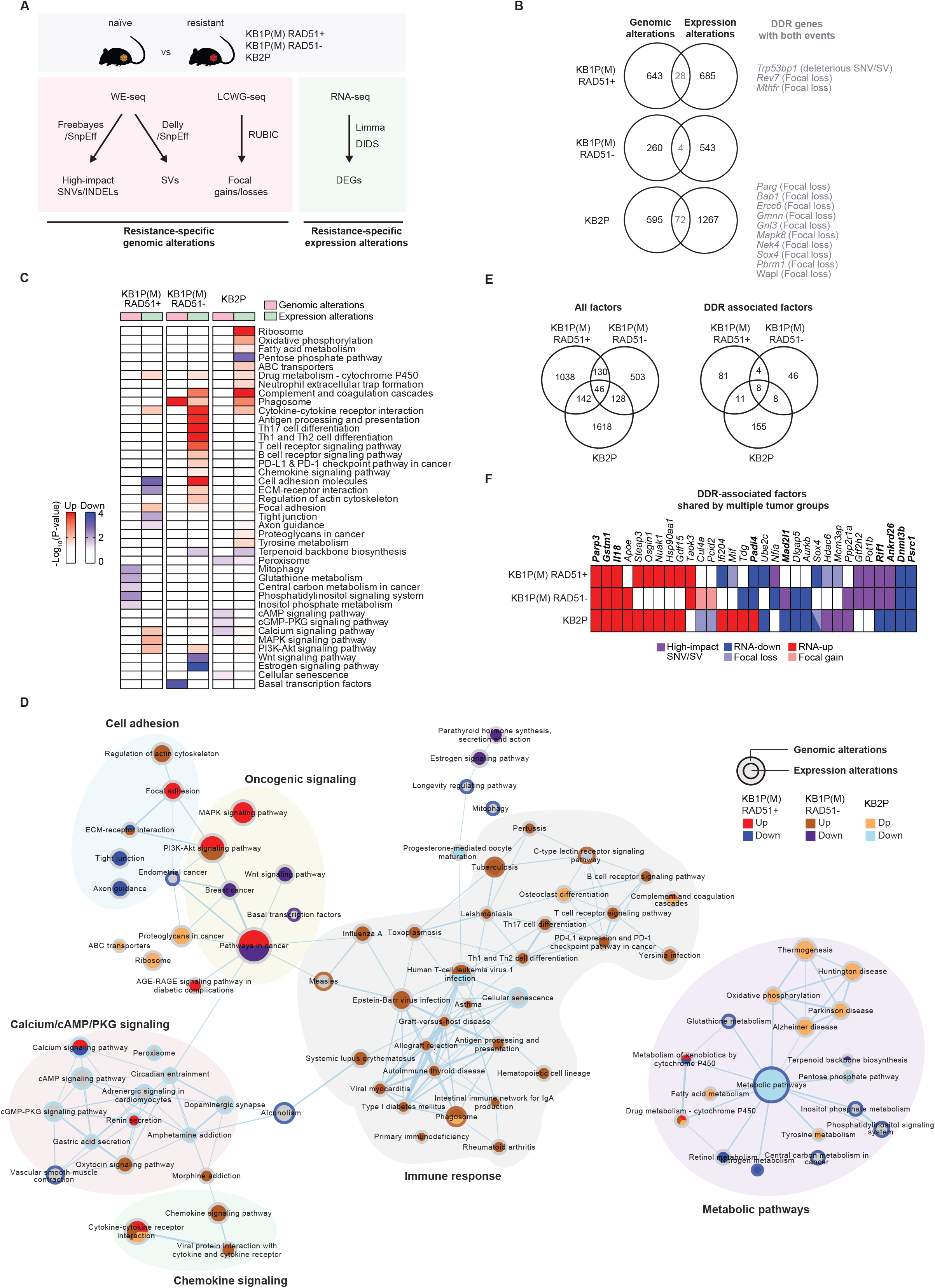
Multi-omics analysis identifies potential PARPi-resistance factors/pathways. **(A)** Schematic overview of the analysis to identify resistance-specific genomic and transcriptional alterations from WE-Seq, LCWG-Seq, and RNA-seq data. For each PARPi naïve and resistant RAD51-positive and RAD51-negative KB1P(M) and KB2P tumors, (1) deleterious SNVs, INDELs, SVs, and focal amplifications/deletions (resistance-specific genomic alterations) and (2) DEG sets identified by either limma or DIDS analysis (resistance-specific transcriptional alterations) were selected. DEG sets were extended to capture the genes with both homogeneous (by limma) and heterogenous behavior (by DIDS) between naïve and resistant tumors. **(B)** Venn diagrams showing the overlaps between the genes having resistance-specific genomic alterations (SNVs, INDELs, SVs, and copy number focal gains and losses) and transcriptional alterations (DEGs) in each resistant tumor type. **(C)** Heatmap of pathways significantly enriched by the genes with resistance-specific genomic and transcriptional alterations. Enrichment, −log_10_(P-value) computed by fisher exact test, by upregulated and downregulated genes are represented by red and green in the heatmap, respectively. **(D)** Pathway enrichment map for genes with resistance-specific genomic and transcriptional alterations constructed by EnrichmentMap (Merico et al., 2010). Node size represents size (number of genes) of gene sets (pathways) and edges represent mutual overlaps between the gene sets (minimum similarity score 0.3). Node and border represent enrichment by the genes with resistance-specific transcriptional and genomic alterations, respectively. **(E)** Venn diagrams showing the overlaps of the genes having resistance-specific alterations with either genomic or transcriptional alterations across the three resistant tumor types. **(F)** List of DDR-associated genes with resistance-specific alterations that were identified in multiple resistant tumor types

To prioritize candidate resistance drivers, we identified genes with genomic alterations that had a significant impact on transcriptional changes in the protein-protein interaction network by carrying out driver analysis with DriverNet (Bashashati et al., 2012). We identified drivers in 25% of genes with resistance-specific mutations and in 23% of genes with copy number alterations (**Figure S4A**). Interestingly, DDR genes were more enriched in drivers compared to non-drivers in all PARPi-resistant tumor groups, with the exception of drivers that were derived from copy number alterations in RAD51-negative KB1P(M) tumors (**Figures S4B-S4D**), supporting the previous findings that DDR pathways are strongly involved in governing PARPi response (Paes Dias et al., 2021).

### *In vitro* loss-of-function screens fail to validate candidate drivers of *in vivo* PARPi resistance

To identify causative drivers among the genes associated with PARPi resistance in KB1P(M) tumors, we performed functional genetic enrichment screens in human BRCA1-deficient cell lines. To obtain a comprehensive set of genes to be functionally tested in the screens, we generated global proteomics and phosphoproteomics data and identified proteins (DE-Prot) and phosphoproteins (DE-Phos) that were differentially expressed between naïve and resistant tumors. We combined the DE-Prot and DE-Phos with genes identified from WE-seq, LCWG-seq and RNA-seq analysis for all KB1P(M) tumors, including tumors whose RAD51-IRIF status was not determined, resulting in 3727 resistance-associated genes (**Figure S4E**). Plausible drivers were prioritized by recurrences in multiple datasets, associations with DDR, and potential as network drivers, yielding a final list of 891 putative PARPi resistance driving factors including 53BP1, REV7, HELB and PARG (**Figure S4F**).

We then generated a focused lentiviral shRNA library targeting the identified 891 candidate genes plus 133 non-essential genes as controls (**Table S5**). Given the strong effect of 53BP1 loss on PARPi resistance, we decided to exclude *Trp53bp1*-targeting shRNAs from the library to prevent them from obscuring less dominant hits. We introduced the lentiviral shRNA library in the human BRCA1-deficient cell lines SUM149PT and RPE1-hTERT-*BRCA1*^-/-^;*TP53*^-/-^, which were subsequently selected with olaparib for three weeks (**Figure S4G**). REV7 behaved as a positive control and was identified as top hit in both cell lines, but no other common hits were found (**Figures S4H and S4I**). We identified RBMS1 as a hit in the screen carried out in SUM149PT cells (**Figure S4H**), but shRNA-mediated depletion of RBMS1 *in vivo* did not significantly affect survival of KB1P tumor-bearing mice (**Figure S4J**). Overall, these results suggest *in vitro* loss-of-function screens are insufficient to validate the candidate drivers of *in vivo* PARPi resistance arising from the multi-omics analysis.

## DISCUSSION

In this study, we combined functional analysis of HR status with multi-omics analysis of a collection of matched PARPi-naïve and PARPi-resistant BRCA1-KO and BRCA2-KO mouse mammary tumors to classify the contribution of previously reported non-reversion resistance mechanisms in a preclinical *“in vivo* reality” of spontaneous acquired resistance. Overall, our analysis highlights the differences in resistance patterns between BRCA1- and BRCA2-deficient tumors and identifies HR restoration via loss of 53BP1 and restoration of PARP signaling via loss of PARG as the two dominant non-reversion resistance mechanisms in BRCA1- and BRCA2-deficient tumors, respectively. Additionally, our analysis generated a catalogue of candidate genes and pathways potentially involved in modulating PARPi response. Expanding the use of PARPi in the clinic should soon provide clinical specimens that will allow us to verify the relevance of our findings.

### Functional differences between BRCA1 and BRCA2 impact PARPi resistance patterns

In our study we also used PARPi resistance as a tool to probe for different activities of BRCA1 and BRCA2 in DNA repair. BRCA1 and BRCA2 are often mentioned together, partly owing to their tumor suppressor activities and roles in HR repair. From a biological standpoint, however, BRCA1 and BRCA2 are not functionally redundant in HR. The epistatic relationship between BRCA1 and BRCA2 was first put forward in the context of embryonic lethality by Ludwig *et al*. almost 20 years ago (Ludwig et al., 1997). Consistent with this relationship, previous work from our laboratory demonstrated that concomitant tissue-specific deletion of the *Brca1* and *Brca2* genes (KB1B2P) resulted in similar tumor development as single gene knockouts (KB1P and KB2P), indicating that BRCA1 and BRCA2 are epistatic in tumor suppression (Liu et al., unpublished data). Our present analysis of PARPi resistance mechanisms reveals a clear functional distinction between BRCA1 and BRCA2 in DNA repair. While HR-deficiency and PARPi sensitivity of BRCA1-KO tumors could be largely suppressed by inactivation of the 53BP1-RIF1-shieldin pathway, BRCA2-KO tumors completely failed to rescue HR activity, as measured by RAD51 foci formation, underlying the essential role of BRCA2 in RAD51 loading during the HR process. Moreover, we found that the levels of the RAD51-IRIF in the PARPi-resistant KB1P(M) tumors were significantly lower than in BRCA-proficient controls, indicating partial restoration of HR activity in these samples (**Figure S1C**). This is consistent with previous DR-GFP-based HR reporter assays we performed in *Tp53bp1-* or *Rev7*-depleted KB1P cells (Xu et al., 2015). In addition, recent studies have suggested that whereas BRCA1 is dispensable for DNA end resection, its interaction with PALB2 and the resulting promotion of RAD51 loading cannot be fully compensated (Belotserkovskaya et al., 2020; Nacson et al., 2018). It is therefore conceivable that inactivation of the 53BP1-RIF1-shieldin pathway in BRCA1-KO tumors rescues the DNA end resection defect, but fails to fully restore HR repair due to lack of BRCA1 activity downstream of resection. Altogether, these data show that BRCA1 and BRCA2 are not epistatic in the repair of DSBs by HR and that these differences impact the resistance patterns in PARPi-treated tumors.

### Resistance mechanisms in HR-restored BRCA1-KO tumors

Loss of 53BP1 was the most frequent alteration in RAD51-positive PARPi-resistant KB1P(M) tumors, whereas all other DSB end-protection factors seemed to be only sporadic. In line with this, mice bearing *Shld1/2-* or *Ctc1*-mutated KB1P tumors exhibit only partial response to olaparib in comparison with mice bearing unmodified KB1P tumors, whereas mice bearing *Trp53bp1*-mutated tumors did not respond to PARPi treatment, resulting in survival curves identical to vehicle-treated mice (Barazas et al., 2018; Noordermeer et al., 2018). Moreover, loss of 53BP1 has also been observed in patient-derived tumor xenograft (PDX) models with acquired resistance to PARPi, and mutations in *TP53BP1* have been reported in tumor biopsies from patients with metastatic BRCA1-associated breast cancer receiving platinum chemotherapy or PARPi (Cruz et al., 2018; Dev et al., 2018; Waks et al., 2020). Altogether, our results indicate that BRCA1-independent HR restoration driven by inactivation of 53BP1 may be the most common mechanism of PARPi resistance in patients with *BRCA1*-mutated tumors that do not undergo BRCA1 reactivation, and that 53BP1 may be a potential biomarker of PARPi response in *BRCA1*-mutated tumors.

### Resistance mechanisms in HR-deficient BRCA1-KO tumors

More than one-third of all PARPi-resistant KB1P(M) tumors were RAD51-negative, indicating that they hadn’t restored HR. Despite the fact that alterations in previously reported resistance factors were found sporadically in PARPi-resistant RAD51-negative KB1P(M) tumors, none of these factors were found significantly altered in any of the analyses carried out in this study. Moreover, alterations in these factors did not occur more frequently in PARPi-resistant RAD51-negative KB1P(M) tumors compared to PARPi-resistant RAD51-positive KB1P(M) or KB2P tumors. Nevertheless, gene set analysis of the DEGs identified in PARPi-resistant RAD51-negative KB1P(M) tumors yielded an enrichment in positive regulation of immune response which was in line with the increase in immune infiltration detected by IHC analysis. In line with this, organoids derived from one of the PARPi-resistant RAD51-negative KB1P(M) tumors (KB1PM7) failed to recapitulate PARPi resistance *in vitro* but upheld PARPi resistance *in vivo*, suggesting PARPi resistance in this tumor may be driven via cell-extrinsic processes that can only be recapitulated *in vivo* (Duarte et al., 2018). Moreover, these tumors preserve PARPi resistance following orthotopic transplantation of tumor fragments into syngeneic mice, indicating that the resistance phenotype is tumor-intrinsic.

Previous studies have reported that *BRCA1/2*-mutated tumors display increased immune infiltration upon treatment with PARPi, suggesting immune infiltration is required for PARPi antitumor efficacy (Pantelidou et al., 2019). Interestingly, among the different pathways upregulated in PARPi-resistant RAD51-negative KB1P(M) tumors, we identified an enrichment in factors associated with upregulation of PD-L1 and PD-1 checkpoint pathway, which is in line with the previously reported upregulation of PD-L1 expression by PARPi (Jiao et al., 2017). PARPi-resistant HR-deficient *BRCA1*-mutated tumors might therefore be treated with combinations of PARPi and immune-checkpoint inhibitors (Ding et al., 2018; Jiao et al., 2017; Shen et al., 2019). Altogether, these data highlight the role of the tumor microenvironment in the response to PARPi.

### Resistance mechanisms in BRCA2-KO tumors

PARG loss was undoubtedly the most frequent alteration in PARPi-resistant KB2P tumors and occurred significantly more often in KB2P tumors than in the other tumor groups, even though a few resistant KB1P(M) tumors carried copy number loss and/or downregulation of *Parg*. The strong selection for PARG loss in KB2P tumors could result from of the impossibility of HR restoration in these tumors. Even so, PARG loss was not more frequent in PARPi-resistant HR-negative KB1P(M) tumors compared to HR-positive KB1P(M) tumors.

Perturbations that occurred more frequently in PARPi-resistant BRCA2-KO tumors than in BRCA1-KO tumors were alterations associated with loss of PARP trapping or rescue of PARP signaling (44% in KB2P versus 32% in KB1P(M); mostly involving *Parg*) and alterations associated with restoration of replication fork stability (56% in KB2P versus 35% in KB1P(M); mostly involving sporadic alterations). Unlike PARG, perturbations in PARP1 were anecdotal in both KB1P and KB2P tumors, suggesting PARP1 activity is critical for the viability of BRCA1/2-deficient cells.

Overall, our data show that PARPi resistance cannot be achieved via HR restoration in BRCA2-deficient tumors that cannot undergo BRCA2 reactivation. Moreover, loss of PARG is the single most recurrent driver of acquired resistance in BRCA2-KO tumors, yielding PARG as a potential predictive marker of PARPi response in patients with *BRCA2*-mutated tumors.

### Limitations of this study

Our tumor collection recapitulates BRCA1/2 loss-driven tumor formation and acquired PARPi resistance but does not capture the full complexity of the human cancer (e.g., metastatic disease, heterogeneity, hypomorphic mutations). Moreover, human genes do not always have mouse orthologs or play exactly the same functions across the two species. For example, the previously reported PARPi resistance factor SLFN11 (Murai et al., 2016) does not have a mouse ortholog and that is why it was not included in our analysis.

This study relies on the comparison with naïve tumors but does not include the comparison with responsive tumors (before developing resistance). Such tumor cohort would allow to identify gene/pathways altered during PARPi treatment before the tumors develop resistance. Additionally, the multi-omics analysis carried out in this study does not include methylome data and thus we might miss epigenetic mechanisms of PARPi resistance. Nevertheless, changes in DNA-methylation are predicted to affect transcriptional activity and should therefore be visible in the RNA-seq data.

In this study we carried out *in vitro* loss-of-function screens in two human cell lines to validate candidate drivers of *in vivo* PARPi resistance, however this approach failed to yield common hits other than the already-known factor *REV7*. A limitation of this approach is that loss-of-function screens, such as shRNA-based screens, can only functionally validate candidate loss-of-function factors, which represent 40% (356/891) of all candidate drivers. Additionally, PARPi resistance in tumors without alterations in known resistance genes could be driven by the additive effect of multiple alterations rather than a single event. Finally, resistance could be driven by genes that modulate the tumor microenvironment, which cannot be adequately modeled in *in vitro* screens.

## ACKNOWLEDGEMENTS

We thank the members of the Preclinical Intervention Unit of the Mouse Clinic for Cancer and Ageing (MCCA) at the NKI for their technical support with the animal studies. We also thank the following NKI core facilities for their excellent service: Animal Pathology facility, Digital Microscopy facility, Genomics Core facility and Animal facility. We would like to thank Piet Borst (NKI, Amsterdam) for scientific discussions. This work was funded by Oncode Institute, the Dutch Cancer Society (KWF 2011-5220 and 2014-6532), the Netherlands Organization for Scientific Research (VICI 91814643), the Netherlands Genomics Initiative (93512009), the Swiss National Science Foundation (310030_179360), the Swiss Cancer League (KLS-4282-08-2017), the Wilhelm-Sander Foundation (no. 2019.069.1) and the European Research Council (ERC-2019-AdG-883877, SyG-319661).

## AUTHOR CONTRIBUTIONS

Conceptualization, J.B., M.P.D., E.G., S.R., J.J.; Methodology, J.B., M.P.D., E.G., F.R., S.R.P.; Investigation, J.B., M.P.D., E.G., F.R., A.A.D., W.S., I.v.d.H, L.B., T.S.K., B.M.; Software Analysis, J.B., E.G., F.R., B.v.d.B., J.R.d.R., R.d.B., R.L.B., C.L.; Original Draft, Review & Editing – J.B., M.P.D., E.G., L.F.A.W., S.R., J.J.; Supervision, M.v.d.V., C.R.J., L.F.A.W., S.R., J.J.; Funding Acquisition, S.R. and J.J.

## RESOURCE AVAILABILITY

### Lead contact

Further information and requests for resources and reagents should be directed to and will be fulfilled by the Lead Contact, Jos Jonkers.

### Materials availability

Materials associated with this study are available upon request from the lead contact.

### Data and code availability

The accession number for the raw data of WE-seq, LCWG-seq and RNA-seq reported in this paper is European Nucleotide Archive (ENA): PRJEB25803.

## EXPERIMENTAL MODEL AND SUBJECT DETAILS

### Cell Lines

KB1P (KB1P-G3) (Jaspers et al., 2013) and KB2P (KB2P-3.4) (Evers et al., 2008) mouse tumor-derived cell lines were previously described and were grown in DMEM/F12 (Gibco) supplemented with 10% FBS and 50 units/ml penicillin-streptomycin (Gibco), containing 5 μg/ml Insulin (Sigma), 5 ng/ml cholera toxin (Sigma) and 5 ng/ml murine epidermal growth-factor (Sigma), under low oxygen conditions (3% O2, 5% CO_2_ at 37°C). RPE1-hTERT *BRCA1^-/-^;TRP53^-/-^* human cell line has been described before (Belotserkovskaya et al., 2020) and was grown in DMEM+GlutaMAX (Gibco) supplemented with 10% FBS and 50 units/ml penicillin-streptomycin (Gibco), under low oxygen conditions (3% O2, 5% CO_2_ at 37°C). SUM149PT (RRID:CVCL_3422) human cell line was grown in RPMI1640 (Gibco) medium supplied with 10% FBS and 50 units/ml penicillin-streptomycin (Gibco), under normal oxygen conditions (21% O2, 5% CO_2_, 37°C).

### Mice

All animal experiments were approved by the Animal Ethics Committee of The Netherlands Cancer Institute (Amsterdam, the Netherlands) and performed in accordance with the Dutch Act on Animal Experimentation (November 2014). Parental FVB (FVB/NRj) and 129/Ola animals were purchased from Janvier Labs and Harlan Olac, respectively, and crossed at the NKI Animal Facility.

## METHOD DETAILS

### *In situ* RAD51-IRIF and 53BP1-IRIF assay

Cryopreserved material of PARPi-naïve or -resistant KB1P, KB1PM and KB2P tumors (KB1P(M): 23 naïve, 47 resistant; KB2P: 19 naïve, 26 resistant) was thawed and orthotopically engrafted into the right mammary fat pad of 6 week-old wild-type syngeneic female mice (KB1P(M) – FVB; KB2P – FVB:129/Ola(F1)). Tumor volume was monitored starting from two weeks after transplantations and calculated using the following formula: 0.5 x length x width^2^. When tumors reached approximately 500 mm^3^ (100% relative tumor volume), they were locally irradiated using a CT-guided high precision cone beam micro-irradiator (X-RAD 225Cx) or left untreated (control). Two different factors were tested to optimize the assay: IR dosage (15 and 24 Gy) and post-irradiation incubation time (1-6 hr). We did not observe significant differences in RAD51 IRIF formation between the two IR dosages, and the highest accumulation of RAD51 foci was detected 2 hours after induction of DNA damage. Based on these results, we analyzed RAD51 IRIF in KB1P(M) and KB2P tumors 2 hr after irradiation with 15Gy. Post irradiation, tumors were isolated and part of the tissue was immediately fixed in 4% (w/v) solution of formaldehyde in PBS (remaining tissue was fresh frozen for the proteomic and phosphoproteomic analyses). 5 μm-thick FFPE (formalin-fixed paraffin embedded) tissue sections were then used for immunofluorescence. Following deparaffinization (70°C, 20 min), tissues were rehydrated and cooked in DAKO Target Retrieval Solution pH 9 (#S236784, DAKO) for 20 min in microwave at ~600W, to allow antigen retrieval. Next, tissue permeabilization was achieved by incubating samples in 0.2% (v/v) Triton X-100 in PBS for 20 min and followed by 1 hr DNAse (1,000 U/ml; #04536282001, Roche) treatment at 37°C. Blocking was done for 30 min in staining buffer (1% (w/v) BSA, 0.15% (w/v) glycine and 0.1% (v/v) Triton X-100 in PBS). Subsequent incubation with primary antibodies was carried out overnight at 4°C, and later with secondary antibodies for 1 hr at room temperature. The following antibodies (diluted in staining buffer) were used in this assay: rabbit polyclonal anti-RAD51 (kind gift from R. Kanaar, Erasmus MC, Rotterdam; 1:5,000), rabbit polyclonal anti-53BP1 (#ab21083, Abcam; 1:1,000), goat polyclonal anti-rabbit, Alexa Fluor^®^ 658-conjugated (#A11011, Thermo Fisher Scientific; diluted 1:1,000). Samples were mounted with VECTASHIELD Hard Set Mounting Media with DAPI (#H-1500; Vector Laboratories). Images were taken with Leica SP5 (Leica Microsystems) confocal system equipped with a x100 objective and image stacks (~6 slices) were analyzed using an in-house developed ImageJ macro to automatically and objectively quantify IR-induced foci, as described before (Xu et al., 2015). Briefly, nuclei were segmented by thresholding the (median-filtered) DAPI signal, followed by a watershed operation to separate touching nuclei. For each *z*-stack the maximum-intensity projection of the foci signal was background-subtracted using a difference of gaussians method. Next, for every nucleus, foci candidates were identified as locations where the resulting pixel values exceeded the background by a factor (typically 25x) times the median standard deviation of all nuclei in the image. Additional filters for discriminating for foci size, nucleus size (to eliminate stromal cells) and absolute brightness were applied. Results were validated by visual inspection. Visualization as well as quantification of foci was done in a blinded fashion. For each sample, five random areas (246 × 246 μm; on average 125 cells per area) were imaged and analyzed. Cell was considered positive if contained >5 nuclear foci. KP tumor was used as a positive control in this assay.

### 53BP1 IHC

All IHC stainings were performed on FFPE material. For 53BP1 IHC, tissue sections were boiled for 30 min in citrate buffer pH 6.0 (#CBB 999, Scytek Laboratories) to facilitate antigen retrieval. Next, the stainings were carried out by using 3% (v/v) H_2_O_2_ solution in methanol for blocking endogenous peroxidase activity (20 min) and 4% BSA plus 5% normal goat serum (NGS) in PBS as a blocking buffer (30 min). Primary antibodies were diluted in 1.25% NGS plus 1% BSA in PBS, and applied on the samples overnight, at 4°C. Incubation with secondary antibodies (diluted in 1.25% NGS/1% BSA in PBS; 30 min, room temperature) (#E0433, DakoCytomation) was followed by incubation with streptavidin conjugated to horseradish peroxidase (1:200; 1.25% NGS/1% BSA in PBS; 30 min) (#P0397, DakoCytomation). For visualization DAB (#D5905, Sigma), H_2_O_2_ (#A-31642, Sigma, 1:1,250) and hematoxylin counterstaining were applied. IHC stainings were evaluated and scored (0 – negative, 1 – low expression, 2 – high expression) by a pathologist who was blinded regarding the identity of the samples. Antibodies used in this protocol: rabbit polyclonal anti-53BP1 (#A300-272A, Bethyl Laboratories; 1:1,000), secondary donkey-anti-goat-Bio (#E0433, DakoCytomation; 1:200), secondary goat-anti-rabbit-Bio (#E0432, Dako Cytomation; 1:1,000).

### Whole-exome sequencing (WE-seq)

Genomic DNA was isolated from fresh frozen tumor tissue using a standard Proteinase K and phenol:chloroform extraction and sheared to approximately 300 bp fragments using Covaris S2 sonicator. Next, 500-1000 ng of sheared DNA was used as a template for a 6-cycle PCR to construct a fragmented library using the KAPA HTP Library Preparation Kit (Roche). Exome enrichment was performed using SeqCap EZ Enrichment Kit (Roche) according to the manufacturer’s protocol (SeqCap EZ Library SR User’s Guide, v5.3). Samples were sequenced on an Illumina HiSeq2500 (Illumina). Adapters in the resulting reads were trimmed using Cutadapt (Martin, 2011) (version 1.12) and the trimmed reads were aligned to the GRCm38 reference genome using BWA (version 0.7.15) (Li, 2013). The resulting alignments were sorted and marked for duplicates with Picard tools (version 2.5.0). Freebayes variant caller was used to identify SNVs and Indels for each sample with the mode of pooled-continuous (min-alternate-fraction = 0.1, min-alternate-count = 3, and min-coverage = 10) (Garrison and Marth, 2012) and resulting variants were annotated by SnpEff (Cingolani et al., 2012). SNVs and Indels identified in FVB/NJ mice were obtained from Sanger Mouse Genome Project (Keane et al., 2011) and used to discard germline variants in our tumor samples. SNVs and Indels that were only identified in resistant tumors and predicted to be high-impact mutations by SnpEff were considered resistant tumor-specific alterations with functional effects and were used for downstream analysis. Structural variants were identified by delly (version 0.9.1) for each pair of resistant and matched naïve tumors and resulting variants were annotated by SnpEff. High-impact SVs predicted by SnpEff were used for downstream analysis.

### Low-coverage whole-genome sequencing (LCWG-seq)

LCWG-seq data was generated as described before (Gogola et al., 2018). Briefly, genomic DNA was isolated from fresh-frozen tumor material using standard phenol:chloroform extraction. LCWG-seq was performed using double stranded DNA (dsDNA) and quantified with the Qubit^®^ dsDNA HS Assay Kit (Invitrogen, #Q32851). Library preparation was performed with 1 μg of DNA and KAPA HTP Library Preparation Kit (KAPA Biosystems, #KK8234). Resulting reads (50 base single-end reads) were trimmed, sorted and marked for duplicates using the same pipeline as for the WES. The resulting alignments were analyzed to generate segmented profile differences between matched (naïve/resistant) samples derived from the same tumor donor using the QDNAseq and QDNAseq.mm10 R packages (bin size = 50K) (Scheinin et al., 2014). To identify regions with recurrent copy number difference (naïve vs resistant), we iteratively ran RUBIC with default cutoff for calling amplifications and deletions (focal threshold = 1e+08, min probes = 4, FDR < 0.25, amp.level and del.level = 0.1) (van Dyk et al., 2016).

### RNA sequencing (RNA-seq)

RNA-seq data was generated as described before (Gogola et al., 2018). Briefly, fresh frozen tumor tissue were placed in 1 ml of TRIsure reagent (#BIO-38032, Bioline) and tissue lysis was achieved by high-speed shaking with stainless steel beads for 10 min, 50 Hz at room temperature (TissueLyser LT, Qiagen). Homogenized tissue lysates were further processed according to the TRIsure manufacturer’s protocol. Strand-specific libraries were generated using the TruSeq Stranded mRNA sample preparation kit (Illumina Inc., San Diego, RS-122-2101/2) according to the manufacturer’s instructions. The resulting reads (50 base single-end reads) were trimmed using Cutadapt (Martin, 2011) and aligned to the GRCm38 reference genome using STAR (version 2.5.2b) (Dobin et al., 2013). To identify differentially expressed (DE) genes, gene expression counts were first generated by featureCounts using gene definitions from Ensembl GRCm38 (version 76) (Liao et al., 2013). Genes with counts per million (CPM) larger than one in at least 10% of samples were used for further analysis. Trimmed mean of M-value (TMM) normalization was applied to the data using edgeR (Robinson et al., 2010) and Limma-voom was used to correct for the donor effect and identify the differentially expressed genes between naïve vs resistant tumors (FDR < 0.25 for KB1P and FDR<0.05 for KB2P, Log2 fold change>0.5) (Law et al., 2014). Because of the intratumoral heterogeneity, we additionally applied DIDS (Detection of Imbalanced Differential Signal) for the detection of subgroup markers in resistant populations (P < 0.05) (de Ronde et al., 2013).

### Selection of previously reported PARPi resistance factors

The list of resistance-associated factors was generated based on previous reports (Paes Dias et al., 2021). We excluded *SLFN11(Murai* et al., 2016) which does not have a mouse ortholog and *Shld3* and *Radx* which were not expressed in our mouse cohort. In total, 25 genes were analyzed in our omics datasets **(Table S1)**.

### Selection of DNA damage response (DDR)-related genes

The DDR gene set was obtained from the previous study that was generated based by merging the gene lists from the previous papers and the NCBI search by terms of “DNA repair”, “DNA damage response”, “DNA replication”, and “telomere-associated genes”(Gogola et al., 2018).

### Gene set over-representation analysis

Gene set over-representation analysis was performed by DAVID (Prakash et al., 2015) for genes with significant upregulation and downregulation (RNA-seq) and focal gains and losses (CNV-seq) in resistant tumors compared to naïve tumors. The significant gene sets were identified as the ones with P<0.05.

### Driver gene analysis

For genes with resistant tumor-specific mutations and copy number variations as mentioned above, DriverNet was used to infer potential driver genes by assessing the impact of alterations on the expression network. Protein-protein interactome (PPI) to construct expression network was obtained by orthologue mapping of human PPI merged from multiple PPI databases (Goel et al.; Hermjakob et al., 2004; Kotlyar et al., 2016; Stark et al., 2006; Xenarios et al., 2000). The genes with more than 1.5-fold changes in expression were defined as genes showing outlying expression and used to assess the impact of mutations in the expression network. P-value was computed by gene-based randomization of 1000 times and genes with P-value <0.05 were selected as potential drivers.

### Mass spectrometry (MS)-based proteomics

For the proteomic and phosphoproteomic analyses of KB1P(M) tumors, we used previously published proteomics dataset generated by MS (PRIDE accession code: PXD032007) (Zingg et al., 2022). For phosphoproteomic analysis, For phosphoproteomic analysis, MaxQuant phosphosite quantification data (Phospho (STY)Sites.txt) was log2-transformed, normalized on the median intensity of all identified phosphosites and replicates averaged favoring data presence. For global protein expression analysis, MaxQuant LFQ Intensity (Cox et al., 2014) was log2-transformed and replicates were averaged. limma (Ritchie et al., 2015) was used for differential expression analysis.

### *Shld2* gene-editing

For CRISPR/Cas9-mediated genome editing of *Shld2*, sgRNA sequences (sgRNA#1: 5’-atcagtcagatccctgcgtt-3’; sgRNA#2: 5’-aacctgagtgatatgactag-3’) were cloned into a modified version of the lentiCRISPR v2 backbone (RRID: Addgene_52961) in which a puromycin resistance ORF was cloned under the hPGK promoter, or into the pX330.puro backbone (Addgene #110403). Cloning of sgRNAs into the lentiCRISPR v2 backbone was carried out by melting the custom DNA oligos (Microsynth) at 95°C for 5 min, followed by annealing at RT for 2h and subsequently ligation with T4 ligase (NEB) into the BsmBI-digested (Fermantas) backbone. Cloning of sgRNAs into the pX330.puro backbone was performed similarly by ligating the previously annealed oligos into the BbsI-HF-digested backbone with T4 ligase (NEB). KB2P tumor-derived cells were transduced with the cloned lentiCRISPR v2 constructs and KB1P cells were transfected with the generated pX330.puro plasmids using a transfection reagent (TransIT-LT1 from Mirus) following the manufacturer’s protocol. All constructs’ sequences were verified by Sanger sequencing.

### Lentiviral Transductions

Lentiviral stocks, pseudotyped with the VSV-G envelope, were generated by transient transfection of HEK293FT cells, as described before (Follenzi et al., 2000). Production of integration-deficient lentivirus (IDLV) stocks was carried out in a similar fashion, with the exception that the packaging plasmid contains a point mutation in the integrase gene (psPAX2, gift from Bastian Evers). Lentiviral titers were determined using the qPCR Lentivirus Titration Kit (Applied Biological Materials), following the manufacturer’s instructions. Cells were incubated with lentiviral supernatants overnight in the presence of polybrene (8 μg/ml). Antibiotic selection was initiated 24h after transduction and was carried out for 3 consecutive days.

### Long-Term Clonogenic Assays

Long-term clonogenic assays were always performed in 6-well plates. KB1P and KB2P tumor-derived cells were seeded at low density to avoid contact inhibition between the clones (4,000 and 3,000 cells/well, respectively). Control untreated plates were fixed with 4% formaldehyde between days 7 and 8 and treated plates between days 8 and 14. For the quantification, cells were stained with 0.1% crystal violet and analyzed in an automated manner using the ImageJ ColonyArea plugin previously described (Guzmán et al., 2014).

### Functional Genetic Enrichment Screens

We have generated a focused shRNA library targeting the human candidate genes plus non-essential genes as controls, resulting in a total of 1025 genes. We selected 5 hairpins per gene, less when 5 weren’t available, resulting in 5062 lentiviral hairpins (pLKO.1) from the Sigma Mission library (TRC 1.0 and 2.0) **(Table S5)**. This library was then used to generate pools of lentiviral shRNAs which were then transduced in RPE1-hTERT *BRCA1^-/-^;TRP53^-/-^* and SUM149PT human cells, as described in the section *Lentiviral Transductions*. Lentiviral transductions were carried out using a multiplicity of transduction (MOI) of 0.3, in order to ensure that each cell only gets incorporated with one only sgRNA. After transduction, the cells stably expressing integrated shRNA were selected with puromycin. After selection, cells were collected (T0) or seeded in the presence of PARPi (SUM149PT:100.000 cells p/ 15 cm plate, 10nM olaparib; RPE1-hTERT *BRCA1^-/-^;TRP53-^/-^:*50.000 cells p/ 15 cm plate, 50nM olaparib). The total number of cells used in a single screen was calculated as following: library complexity x coverage (1000x). Triplicates were carried out for both cell lines. Cells were kept in culture for 3 weeks and medium was refreshed every 5 days. In the end of the screen, or at T0, cells were pooled and genomic DNA was extracted (QIAmp DNA Mini Kit, Qiagen). shRNA sequences were retrieved by a two-step PCR amplification, as described before (Xu et al., 2015). To maintain screening coverage, the amount of genomic DNA used as an input for the first PCR reaction was taken into account (6 μg of genomic DNA per 10^6^ genomes, 1 μg/PCR reaction). Resulting PCR products were purified using MiniElute PCR Purification Kit (Qiagen) and submitted for Illumina sequencing. Sequence alignment and dropout analysis was carried out using the algorithm MaGECK (Li et al., 2014).

### Statistical analysis

High-throughput genomics, transcriptomics, proteomics and phosphoproteomics data were analyzed as described in relevant Method sections. Statistical analysis of long-term clonogenic assays was performed using 2-way ANOVA followed by Dunnett’s test. For the analysis of RAD51/53BP1 IRIF data we used two-tailed Mann-Whitney *U* test. **** p < 0.0001, *** p < 0.001, ** p < 0.01.

## REFERENCES

Ang, J.E., Gourley, C., Powell, C.B., High, H., Shapira-Frommer, R., Castonguay, V., De Greve, J., Atkinson, T., Yap, T.A., Sandhu, S., et al. (2013). Efficacy of chemotherapy in BRCA1/2 mutation carrier ovarian cancer in the setting of PARP inhibitor resistance: a multi-institutional study. Clin. Cancer Res. 19, 5485–5493.

Barazas, M., Annunziato, S., Pettitt, S.J., de Krijger, I., Ghezraoui, H., Roobol, S.J., Lutz, C., Frankum, J., Song, F.F., Brough, R., et al. (2018). The CST Complex Mediates End Protection at Double-Strand Breaks and Promotes PARP Inhibitor Sensitivity in BRCA1-Deficient Cells. Cell Rep. 23, 2107–2118.

Barber, L.J., Sandhu, S., Chen, L., Campbell, J., Kozarewa, I., Fenwick, K., Assiotis, I., Rodrigues, D.N., Reis Filho, J.S., Moreno, V., et al. (2013). Secondary mutations in BRCA2 associated with clinical resistance to a PARP inhibitor. J. Pathol. 229, 422–429.

Bashashati, A., Haffari, G., Ding, J., Ha, G., Lui, K., Rosner, J., Huntsman, D.G., Caldas, C., Aparicio, S.A., and Shah, S.P. (2012). DriverNet: uncovering the impact of somatic driver mutations on transcriptional networks in cancer. Genome Biol. 13, R124–R124.

Belotserkovskaya, R., Raga Gil, E., Lawrence, N., Butler, R., Clifford, G., Wilson, M.D., and Jackson, S.P. (2020). PALB2 chromatin recruitment restores homologous recombination in BRCA1-deficient cells depleted of 53BP1. Nat. Commun. 11, 1–11.

Bhattacharya, A., Bense, R.D., Urzúa-Traslaviña, C.G., de Vries, E.G.E., van Vugt, M.A.T.M., and Fehrmann, R.S.N. (2020). Transcriptional effects of copy number alterations in a large set of human cancers. Nat. Commun. 11, 1–12.

Boersma, V., Moatti, N., Segura-Bayona, S., Peuscher, M.H., van der Torre, J., Wevers, B. a, Orthwein, A., Durocher, D., and Jacobs, J.J.L. (2015). MAD2L2 controls DNA repair at telomeres and DNA breaks by inhibiting 5’ end resection. Nature 521, 537–540.

Bouwman, P., Aly, A., Escandell, J.M., Pieterse, M., Bartkova, J., van der Gulden, H., Hiddingh, S., Thanasoula, M., Kulkarni, A., Yang, Q., et al. (2010). 53BP1 loss rescues BRCA1 deficiency and is associated with triple-negative and BRCA-mutated breast cancers. Nat. Struct. Mol. Biol. 17, 688–695.

Bryant, H.E., Schultz, N., Thomas, H.D., Parker, K.M., Flower, D., Lopez, E., Kyle, S., Meuth, M., Curtin, N.J., and Helleday, T. (2005). Specific killing of BRCA2-deficient tumours with inhibitors of poly(ADP-ribose) polymerase. Nature 434, 913–917.

Bunting, S.F., Callén, E., Wong, N., Chen, H.-T., Polato, F., Gunn, A., Bothmer, A., Feldhahn, N., Fernandez-Capetillo, O., Cao, L., et al. (2010). 53BP1 Inhibits Homologous Recombination in Brca1-Deficient Cells by Blocking Resection of DNA Breaks. Cell 141, 243–254.

Cantor, S.B., and Calvo, J.A. (2017). Fork Protection and Therapy Resistance in Hereditary Breast Cancer.

Chapman, J.R., Barral, P., Vannier, J.B., Borel, V., Steger, M., Tomas-Loba, A., Sartori, A.A., Adams, I.R., Batista, F.D., and Boulton, S.J. (2013). RIF1 Is Essential for 53BP1-Dependent Nonhomologous End Joining and Suppression of DNA Double-Strand Break Resection. Mol. Cell 49, 858–871.

Cingolani, P., Platts, A., Wang, L.L., Coon, M., Nguyen, T., Wang, L., Land, S.J., Lu, X., and Ruden, D.M. (2012). A program for annotating and predicting the effects of single nucleotide polymorphisms, SnpEff: SNPs in the genome of Drosophila melanogaster strain w1118; iso-2; iso-3. Fly (Austin). 6, 80–92.

Cox, J., Hein, M.Y., Luber, C.A., Paron, I., Nagaraj, N., and Mann, M. (2014). Accurate proteome-wide label-free quantification by delayed normalization and maximal peptide ratio extraction, termed MaxLFQ. Mol. Cell. Proteomics 13, 2513–2526.

Cruz, C., Castroviejo-Bermejo, M., Gutiérrez-Enríquez, S., Llop-Guevara, A., Ibrahim, Y.H., Gris-Oliver, A., Bonache, S., Morancho, B., Bruna, A., Rueda, O.M., et al. (2018). RAD51 foci as a functional biomarker of homologous recombination repair and PARP inhibitor resistance in germline BRCA-mutated breast cancer. Ann. Oncol. 29, 1203–1210.

Dev, H., Chiang, T.-W.W., Lescale, C., de Krijger, I., Martin, A.G., Pilger, D., Coates, J., Sczaniecka-Clift, M., Wei, W., Ostermaier, M., et al. (2018). Shieldin complex promotes DNA end-joining and counters homologous recombination in BRCA1-null cells. Nat. Cell Biol. 20, 954–965.

Ding, L., Kim, H.J., Wang, Q., Kearns, M., Jiang, T., Ohlson, C.E., Li, B.B., Xie, S., Liu, J.F., Stover, E.H., et al. (2018). PARP Inhibition Elicits STING-Dependent Antitumor Immunity in Brca1-Deficient Ovarian Cancer. Cell Rep. 25, 2972–2980.e5.

Dobin, A., Davis, C.A., Schlesinger, F., Drenkow, J., Zaleski, C., Jha, S., Batut, P., Chaisson, M., and Gingeras, T.R. (2013). STAR: ultrafast universal RNA-seq aligner. Bioinformatics 29, 15–21.

Domchek, S.M. (2017). Reversion Mutations with Clinical Use of PARP Inhibitors: Many Genes, Many Versions. Cancer Discov. 7, 937–939.

Duarte, A.A., Gogola, E., Sachs, N., Barazas, M., Annunziato, S., R de Ruiter, J., Velds, A., Blatter, S., Houthuijzen, J.M., van de Ven, M., et al. (2018). BRCA-deficient mouse mammary tumor organoids to study cancer-drug resistance. Nat. Methods 15, 134–140.

van Dyk, E., Hoogstraat, M., Ten Hoeve, J., Reinders, M.J.T., and Wessels, L.F.A. (2016). RUBIC identifies driver genes by detecting recurrent DNA copy number breaks. Nat. Commun. 7, 12159.

Escribano-Díaz, C., Orthwein, A., Fradet-Turcotte, A., Xing, M., Young, J.T.F., Tkác, J., Cook, M.A., Rosebrock, A.P., Munro, M., Canny, M.D., et al. (2013). A cell cycle-dependent regulatory circuit composed of 53BP1-RIF1 and BRCA1-CtIP controls DNA repair pathway choice. Mol. Cell 49, 872–883.

Evers, B., Drost, R., Schut, E., de Bruin, M., van der Burg, E., Derksen, P.W.B., Holstege, H., Liu, X., van Drunen, E., Beverloo, H.B., et al. (2008). Selective Inhibition of BRCA2-Deficient Mammary Tumor Cell Growth by AZD2281 and Cisplatin. Clin. Cancer Res. 14, 3916–3925.

Farmer, H., Nuala, M., J. Lord, C., N. J. Tutt, A., A. Johnson, D., B. Richardson, T., and Santarosa, M. (2005). Targeting the DNA repair defect in BRCA mutant cells as a therapeutic strategy. Nature 434, 917–921.

Feng, L., Li, N., Li, Y., Wang, J., Gao, M., Wang, W., and Chen, J. (2015). Cell cycle-dependent inhibition of 53BP1 signaling by BRCA1. Cell Discov. 1, 15019.

Findlay, S., Heath, J., Luo, V.M., Malina, A., Morin, T., Coulombe, Y., Djerir, B., Li, Z., Samiei, A., Simo-Cheyou, E., et al. (2018). SHLD 2/ FAM 35A co-operates with REV 7 to coordinate DNA double-strand break repair pathway choice. EMBO J. 37.

Follenzi, A., Ailles, L.E., Bakovic, S., Geuna, M., and Naldini, L. (2000). Gene transfer by lentiviral vectors is limited by nuclear translocation and rescued by HIV-1 pol sequences. Nat. Genet. 25, 217–222.

Ganesan, S. (2018). Tumor Suppressor Tolerance: Reversion Mutations in BRCA1 and BRCA2 and Resistance to PARP Inhibitors and Platinum. JCO Precis. Oncol. 1–4.

Gao, S., Feng, S., Ning, S., Liu, J., Zhao, H., Xu, Y., Shang, J., Li, K., Li, Q., Guo, R., et al. (2018). An OB-fold complex controls the repair pathways for DNA double-strand breaks. Nat. Commun. 9, 1–10.

Garrison, E., and Marth, G. (2012). Haplotype-based variant detection from short-read sequencing.

Ghezraoui, H., Oliveira, C., Becker, J.R., Bilham, K., Moralli, D., Anzilotti, C., Fischer, R., Deobagkar-Lele, M., Sanchiz-Calvo, M., Fueyo-Marcos, E., et al. (2018). 53BP1 cooperation with the REV7–shieldin complex underpins DNA structure-specific NHEJ. Nature 560, 122–127.

Godin, S.K., Sullivan, M.R., and Bernstein, K.A. (2016). Novel insights into RAD51 activity and regulation during homologous recombination and DNA replication. Biochem. Cell Biol. 94, 407–418.

Goel, R., Harsha, H.C., Pandey, A., and Keshava Prasad, T.S. Human Protein Reference Database and Human Proteinpedia as Resources for Phosphoproteome Analysis.

Gogola, E., Duarte, A.A., de Ruiter, J.R., Wiegant, W.W., Schmid, J.A., de Bruijn, R., James, D.I., Guerrero Llobet, S., Vis, D.J., Annunziato, S., et al. (2018). Selective Loss of PARG Restores PARylation and Counteracts PARP Inhibitor-Mediated Synthetic Lethality. Cancer Cell 33, 1078–1093.e12.

Gupta, R., Somyajit, K., Narita, T., Maskey, E., Stanlie, A., Kremer, M., Typas, D., Lammers, M., Mailand, N., Nussenzweig, A., et al. (2018). DNA Repair Network Analysis Reveals Shieldin as a Key Regulator of NHEJ and PARP Inhibitor Sensitivity. Cell 173, 972–988.e23.

Guzmán, C., Bagga, M., Kaur, A., Westermarck, J., and Abankwa, D. (2014). ColonyArea: an ImageJ plugin to automatically quantify colony formation in clonogenic assays. PLoS One 9.

Hermjakob, H., Montecchi-Palazzi, L., Lewington, C., Mudali, S., Kerrien, S., Orchard, S., Vingron, M., Roechert, B., Roepstorff, P., Valencia, A., et al. (2004). IntAct: an open source molecular interaction database. Nucleic Acids Res. 32, D452.

Jaspers, J.E., Kersbergen, A., Boon, U., Sol, W., Van Deemter, L., Zander, S.A., Drost, R., Wientjens, E., Ji, J., Aly, A., et al. (2013). Loss of 53BP1 causes PARP inhibitor resistance in BRCA1-mutated mouse mammary tumors. Cancer Discov. 3, 68–81.

Jiao, S., Xia, W., Yamaguchi, H., Wei, Y., Chen, M.K., Hsu, J.M., Hsu, J.L., Yu, W.H., Du, Y., Lee, H.H., et al. (2017). PARP Inhibitor Upregulates PD-L1 Expression and Enhances Cancer-Associated Immunosuppression. Clin. Cancer Res. 23, 3711–3720.

Jonkers, J., Meuwissen, R., van der Gulden, H., Peterse, H., van der Valk, M., and Berns, A. (2001). Synergistic tumor suppressor activity of BRCA2 and p53 in a conditional mouse model for breast cancer. Nat. Genet. 29, 418–425.

Kas, S.M., De Ruiter, J.R., Schipper, K., Schut, E., Bombardelli, L., Wientjens, E., Drenth, A.P., De Korte-Grimmerink, R., Mahakena, S., Phillips, C., et al. (2018). Transcriptomics and Transposon Mutagenesis Identify Multiple Mechanisms of Resistance to the FGFR Inhibitor AZD4547. Cancer Res. 78, 5668–5679.

Keane, T.M., Goodstadt, L., Danecek, P., White, M.A., Wong, K., Yalcin, B., Heger, A., Agam, A., Slater, G., Goodson, M., et al. (2011). Mouse genomic variation and its effect on phenotypes and gene regulation. Nat. 2011 4777364 477, 289–294.

Kotlyar, M., Pastrello, C., Sheahan, N., and Jurisica, I. (2016). Integrated interactions database: tissue-specific view of the human and model organism interactomes. Nucleic Acids Res. 44, D536–D541.

Law, C.W., Chen, Y., Shi, W., and Smyth, G.K. (2014). voom: Precision weights unlock linear model analysis tools for RNA-seq read counts. Genome Biol. 15, R29–R29.

Li, H. (2013). Aligning sequence reads, clone sequences and assembly contigs with BWA-MEM.

Liao, Y., Smyth, G.K., and Shi, W. (2013). featureCounts: an efficient general purpose program for assigning sequence reads to genomic features. Bioinformatics 30, 923–930.

Litton, J.K., Scoggins, M.E., Hess, K.R., Adrada, B.E., Murthy, R.K., Damodaran, S., DeSnyder, S.M., Brewster, A.M., Barcenas, C.H., Valero, V., et al. (2020). Neoadjuvant Talazoparib for Patients With Operable Breast Cancer With a Germline BRCA Pathogenic Variant. J. Clin. Oncol. 38, 388–394.

Liu, X., Holstege, H., van der Gulden, H., Treur-Mulder, M., Zevenhoven, J., Velds, A., Kerkhoven, R.M., van Vliet, M.H., Wessels, L.F.A., Peterse, J.L., et al. (2007). Somatic loss of BRCA1 and p53 in mice induces mammary tumors with features of human BRCA1-mutated basal-like breast cancer. Proc. Natl. Acad. Sci. 104, 12111–12116.

Ludwig, T., Chapman, D.L., Papaioannou, V.E., and Efstratiadis, A. (1997). Targeted mutations of breast cancer susceptibility gene homologs in mice: lethal phenotypes of Brca1, Brca2, Brca1/Brca2, Brca1/p53, and Brca2/p53 nullizygous embryos. Genes Dev. 11, 1226–1241.

Lupo, B., and Trusolino, L. (2014). Inhibition of poly(ADP-ribosyl)ation in cancer: Old and new paradigms revisited. Biochim. Biophys. Acta - Rev. Cancer 1846, 201–215.

Martin, M. (2011). Cutadapt removes adapter sequences from high-throughput sequencing reads. EMBnet.Journal 17, 10.

Merico, D., Isserlin, R., Stueker, O., Emili, A., and Bader, G.D. (2010). Enrichment Map: A Network-Based Method for Gene-Set Enrichment Visualization and Interpretation. PLoS One 5, e13984.

Murai, J., Huang, S.Y.N., Das, B.B., Renaud, A., Zhang, Y., Doroshow, J.H., Ji, J., Takeda, S., and Pommier, Y. (2012). Trapping of PARP1 and PARP2 by clinical PARP inhibitors. Cancer Res. 72, 5588–5599.

Murai, J., Feng, Y., Yu, G.K., Ru, Y., Tang, S.-W., Shen, Y., and Pommier, Y. (2016). Resistance to PARP inhibitors by SLFN11 inactivation can be overcome by ATR inhibition. Oncotarget 7, 76534–76550.

Murai, J., Tang, S.-W., Leo, E., Baechler, S.A., Redon, C.E., Zhang, H., Al Abo, M., Rajapakse, V.N., Nakamura, E., Jenkins, L.M.M., et al. (2018). SLFN11 Blocks Stressed Replication Forks Independently of ATR. Mol. Cell 69, 371–384.e6.

Nacson, J., Krais, J.J., Bernhardy, A.J., Clausen, E., Feng, W., Wang, Y., Nicolas, E., Cai, K.Q., Tricarico, R., Hua, X., et al. (2018). BRCA1 Mutation-Specific Responses to 53BP1 Loss-Induced Homologous Recombination and PARP Inhibitor Resistance. Cell Rep. 24, 3513–3527.e7.

Noordermeer, S.M., Adam, S., Setiaputra, D., Barazas, M., Pettitt, S.J., Ling, A.K., Olivieri, M., Álvarez-Quilón, A., Moatti, N., Zimmermann, M., et al. (2018). The shieldin complex mediates 53BP1-dependent DNA repair. Nature 560, 117–121.

Norquist, B., Wurz, K.A., Pennil, C.C., Garcia, R., Gross, J., Sakai, W., Karlan, B.Y., Taniguchi, T., and Swisher, E.M. (2011). Secondary somatic mutations restoring BRCA1/2 predict chemotherapy resistance in hereditary ovarian carcinomas. J. Clin. Oncol. 29, 3008–3015.

Oplustil O’Connor, L., Rulten, S.L., Cranston, A.N., Odedra, R., Brown, H., Jaspers, J.E., Jones, L., Knights, C., Evers, B., Ting, A., et al. (2016). The PARP Inhibitor AZD2461 Provides Insights into the Role of PARP3 Inhibition for Both Synthetic Lethality and Tolerability with Chemotherapy in Preclinical Models. Cancer Res. 76, 6084–6094.

Paes Dias, M., Moser, S.C., Ganesan, S., and Jonkers, J. (2021). Understanding and overcoming resistance to PARP inhibitors in cancer therapy. Nat. Rev. Clin. Oncol. 1–19.

Pantelidou, C., Sonzogni, O., Taveira, M.D.O., Mehta, A.K., Kothari, A., Wang, D., Visal, T., Li, M.K., Pinto, J., Castrillon, J.A., et al. (2019). PARP Inhibitor Efficacy Depends on CD8 + T-cell Recruitment via Intratumoral STING Pathway Activation in BRCA-Deficient Models of Triple-Negative Breast Cancer. Cancer Discov. 9, 722–737.

Pettitt, S.J., Rehman, F.L., Bajrami, I., Brough, R., Wallberg, F., Kozarewa, I., Fenwick, K., Assiotis, I., Chen, L., Campbell, J., et al. (2013). A genetic screen using the PiggyBac transposon in haploid cells identifies Parp1 as a mediator of olaparib toxicity. PLoS One 8, e61520.

Pettitt, S.J., Krastev, D.B., Brandsma, I., Dréan, A., Song, F., Aleksandrov, R., Harrell, M.I., Menon, M., Brough, R., Campbell, J., et al. (2018). Genome-wide and high-density CRISPR-Cas9 screens identify point mutations in PARP1 causing PARP inhibitor resistance. Nat. Commun. 9, 1849.

Prakash, R., Zhang, Y., Feng, W., and Jasin, M. (2015). Homologous recombination and human health: The roles of BRCA1, BRCA2, and associated proteins. Cold Spring Harb. Perspect. Biol. 7.

Ray Chaudhuri, A., and Nussenzweig, A. (2017). The multifaceted roles of PARP1 in DNA repair and chromatin remodelling.

Ray Chaudhuri, A., Callen, E., Ding, X., Gogola, E., Duarte, A.A., Lee, J.-E., Wong, N., Lafarga, V., Calvo, J.A., Panzarino, N.J., et al. (2016). Replication fork stability confers chemoresistance in BRCA-deficient cells. Nature 535, 382–387.

Ritchie, M.E., Phipson, B., Wu, D., Hu, Y., Law, C.W., Shi, W., and Smyth, G.K. (2015). limma powers differential expression analyses for RNA-sequencing and microarray studies. Nucleic Acids Res. 43, e47–e47.

Robinson, M.D., McCarthy, D.J., and Smyth, G.K. (2010). edgeR: a Bioconductor package for differential expression analysis of digital gene expression data. Bioinformatics 26, 139–140.

de Ronde, J.J., Rigaill, G., Rottenberg, S., Rodenhuis, S., and Wessels, L.F.A. (2013). Identifying subgroup markers in heterogeneous populations. Nucleic Acids Res. 41, e200–e200.

Rondinelli, B., Gogola, E., Yücel, H., Duarte, A.A., Van De Ven, M., Van Der Sluijs, R., Konstantinopoulos, P.A., Jonkers, J., Ceccaldi, R., Rottenberg, S., et al. (2017). EZH2 promotes degradation of stalled replication forks by recruiting MUS81 through histone H3 trimethylation. Nat. Cell Biol. 19, 1371–1378.

Rottenberg, S., Nygren, A.O.H., Pajic, M., van Leeuwen, F.W.B., van der Heijden, I., van de Wetering, K., Liu, X., de Visser, K.E., Gilhuijs, K.G., van Tellingen, O., et al. (2007). Selective induction of chemotherapy resistance of mammary tumors in a conditional mouse model for hereditary breast cancer. Proc. Natl. Acad. Sci. U. S. A. 104, 12117–12122.

Scheinin, I., Sie, D., Bengtsson, H., van de Wiel, M.A., Olshen, A.B., van Thuijl, H.F., van Essen, H.F., Eijk, P.P., Rustenburg, F., Meijer, G.A., et al. (2014). DNA copy number analysis of fresh and formalin-fixed specimens by shallow whole-genome sequencing with identification and exclusion of problematic regions in the genome assembly. Genome Res. 24, 2022–2032.

Scully, R., Panday, A., Elango, R., and Willis, N.A. (2019). DNA double-strand break repair-pathway choice in somatic mammalian cells. Nat. Rev. Mol. Cell Biol. 20, 698–714.

Shen, J., Zhao, W., Ju, Z., Wang, L., Peng, Y., Labrie, M., Yap, T.A., Mills, G.B., and Peng, G. (2019). PARPi Triggers the STING-Dependent Immune Response and Enhances the Therapeutic Efficacy of Immune Checkpoint Blockade Independent of BRCAness. Cancer Res. 79, 311–319.

Stark, C., Breitkreutz, B.J., Reguly, T., Boucher, L., Breitkreutz, A., and Tyers, M. (2006). BioGRID: a general repository for interaction datasets. Nucleic Acids Res. 34.

Szklarczyk, D., Gable, A.L., Nastou, K.C., Lyon, D., Kirsch, R., Pyysalo, S., Doncheva, N.T., Legeay, M., Fang, T., Bork, P., et al. (2021). The STRING database in 2021: customizable protein-protein networks, and functional characterization of user-uploaded gene/measurement sets. Nucleic Acids Res. 49, D605–D612.

Tomida, J., Takata, K., Bhetawal, S., Person, M.D., Chao, H., Tang, D.G., and Wood, R.D. (2018). FAM 35A associates with REV 7 and modulates DNA damage responses of normal and BRCA 1-defective cells. EMBO J. 37, e99543.

Di Virgilio, M., Callen, E., Yamane, A., Zhang, W., Jankovic, M., Gitlin, A.D., Feldhahn, N., Resch, W., Oliveira, T.Y., Chait, B.T., et al. (2013). Rif1 prevents resection of DNA breaks and promotes immunoglobulin class switching. Science (80-.). 339, 711–715.

Waks, A.G., Cohen, O., Kochupurakkal, B., Kim, D., Dunn, C.E., Buendia Buendia, J., Wander, S., Helvie, K., Lloyd, M.R., Marini, L., et al. (2020). Reversion and non-reversion mechanisms of resistance to PARP inhibitor or platinum chemotherapy in BRCA1/2-mutant metastatic breast cancer. Ann. Oncol. 31, 590–598.

Weigelt, B., Comino-Méndez, I., de Bruijn, I., Tian, L., Meisel, J.L., García-Murillas, I., Fribbens, C., Cutts, R., Martelotto, L.G., Ng, C.K.Y., et al. (2017). Diverse BRCA1 and BRCA2 Reversion Mutations in Circulating Cell-Free DNA of Therapy-Resistant Breast or Ovarian Cancer. Clin. Cancer Res. 23, 6708–6720.

Xenarios, I., Rice, D.W., Salwinski, L., Baron, M.K., Marcotte, E.M., and Eisenberg, D. (2000). DIP: the Database of Interacting Proteins. Nucleic Acids Res. 28, 289.

Xu, G., Ross Chapman, J., Brandsma, I., Yuan, J., Mistrik, M., Bouwman, P., Bartkova, J., Gogola, E., Warmerdam, D., Barazas, M., et al. (2015). REV7 counteracts DNA double-strand break resection and affects PARP inhibition. Nature 521, 541–544.

Zimmermann, M., Lottersberger, F., Buonomo, S.B., Sfeir, A., and de Lange, T. (2013). 53BP1 Regulates DSB Repair Using Rif1 to Control 5’ End Resection. Science (80-.). 339, 700–704.

Zingg, D., Bhin, J., Yemelyanenko, J., Kas, S.M., Rolfs, F., Lutz, C., Lee, J.K., Klarenbeek, S., Silverman, I.M., Annunziato, S., et al. (2022). Truncated FGFR2 is a clinically actionable oncogene in multiple cancers. Nat. 2022 1–9.

