## Supplemental Figure legends for "Multi-omics analysis reveals distinct non-reversion mechanisms of PARPi resistance in BRCA1- versus BRCA2-deficient mammary tumors"

**SUPLEMENTARY FIGURE LEGENDS**

**Figure S1.** **Optimization of the RAD51 IRIF formation assay. Related to Figure 1**

**(A)** Percentage of RAD51 positive cells (>5 foci/nucleus) per tumor area (single data point) in a KP tumor irradiated with 0, 15 or 24 Gy, 1-6 hr post-irradiation; ****p<0.0001, n.s. not significant (two-tailed Mann-Whitney U test); data shown as mean (red line) ± SD of a replicate, the experiment was repeated twice; grey dotted line indicates the mean value of a sample irradiated with 15 Gy and incubated for 2 hr.

**(B)** Number of foci per nucleus (single data point) quantified for the total tumor cell population of a KP described in (A); mean values are represented by green lines; **p<0.01, statistical analysis as in (A).

**(C)** Number of foci per nucleus quantified for positive cell population (>5 foci/nucleus) of a KP tumor described in (A), and represented as violin plots showing the density (width = frequency) of the data; mean values are represented by blue dots; statistical analysis as in (A).

**(D)** Summary of analyses represented in (A-C).

**(E-F)** Growth curves of KB1P(M) (E) and KB2P (F) tumors, samples were irradiated at day 7.

**(G-H)** Visualization of exome sequencing reads for the *Brca1* (G) and *Brca2* (H) genes in KB1P(M) and KB2P tumor samples, respectively; and normal tissues (spleen 1/2 and liver 1/2; controls), showing that deletions of specific exons (marked in red; *Brca1* – exons 4-12 (GRCm38, previously referred to as exons 5-13 (Liu et al., 2007), *Brca2* – exon 11) are preserved in PARPi-naïve and PARPi-resistant tumor samples; data representative for the whole KB1P(M) and KB2P tumor panels.

**Figure S2. *Trp53bp1* gene expression correlates with 53BP1 IRIF. Related to Figure 2**

**(A-B)** Representative images (left) and quantification (right) of clonogenic assays in the presence of olaparib in KB1P (A) and KB2P (B) tumor-derived cells modified by CRISPR-Cas9 with the indicated sgRNAs. Control untreated plates were harvested on day 7/8 and treated plates on day 14. Statistical analysis was performed using 2-way ANOVA followed by Dunnett’s test. ∗p < 0.05, ∗∗p < 0.01, ∗∗∗p < 0.001.

**(C-D)** Representative images (A) and quantification (B) of 53BP1 IRIFs for the different matched KB1P(M) tumor pairs; IR – irradiated, NIR – non irradiated; scale bar, 100 µm; data in (B) represented as percentage of positive cells (≥5 foci/nucleus) per imaged area (single data point, typically 100-200 cells/area); ****p<0.0001, ***p<0.001, **p<0.01, n.s. not significant (two-tailed Mann-Whitney U test).

**(E)** Outcome of the RAD51 and 53BP1 IRIF assays for PARPi-resistant KB2P tumors and for PARPi-naïve KB1P(M) and KB2P tumors; N – naïve; R – resistant.

**Figure S3. Systematic analysis of genomic and transcriptomics data for PARP-naïve vs PARPi-resistant KB1P(M) and KB2P tumors. Related to Figure 3**

**(A-C)** Identification of focal gains and losses in RAD51-positive (A), RAD51-negative (B) KB1P(M) resistant tumors, and KB2P resistant tumors (C) compared to matched naïve tumors using RUBIC (van Dyk et al., 2016). Genes were highlighted if they are previously reported PARPi-resistance factors.

**(D)** Co-functionality network of DDR genes with significant focal gain or loss in RAD51-positive KB1P(M) (black), RAD51-negative KB1P(M) (gray) and KB2P (brown) resistant tumors. Gain (red) or loss (blue) are represented by border color DDR, DNA damage response. Top 10 pathways enriched by the genes in the network are listed with Z-transformed P value. The network construction and pathway analysis were performed by the GenetICA-Network (Bhattacharya et al., 2020).

**(E-G)** Gene set analysis represented by the genes with significant focal gain or loss in PARPi-resistant RAD51-positive KB1P(M) (E), RAD51-negative KB1P(M) (F) and KB2P (G) tumors, detected by RUBIC. Because only focal losses were detected in RAD51-positive KB1P(M) resistant tumors, gene set analysis was performed for the genes with focal losses (blue). For RAD51-negative KB1P(M) resistant tumors, only one focal gain was detected, gene set analysis was therefore performed only for the genes with focal gain (red). KB2P tumors showed three focal losses and one focal gain. The focal gain only encoded mostly for non-protein-coding genes with the exception of three genes and thus gene set analysis was performed only for the genes encoded by the areas of focal losses. The gene sets with P<0.01 (fisher’s exact test) were presented.

**(H-J)** Volcano plots of differentially expressed genes (DEGs) in PARPi-naïve vs PARPi-resistant RAD51-positive KB1P(M) (H), RAD51-negative KB1P(M) (I) and KB2P (J) tumors. Genes were highlighted if they are previously reported PARPi-resistance factors.

**Figure S4. Loss-of-function genetic screens are insufficient to validate the different dimensions of our findings. Related to Figure 4.**

**(A)** Bar plot representing the number of driver and non-driver genes with high impact on expression of neighbors in a protein-protein interaction network. Driver potential was assessed for the genes with resistance-specific mutations (left) and copy number variations (right) by DriverNet (Bashashati et al., 2012).

**(B)** Enrichment of DDR genes in the identified drivers from each resistant tumor group. Log_2_ odd ratios (proportion of DDR genes in driver genes vs non-driver genes) were presented with 95% confidence intervals. P-values were computed by fisher’s exact test.

**(C-D)** DDR genes identified as drivers from resistance-associated mutations (C) and CNVs (D) were presented in the co-functionality networks.

**(E)** Schematic diagram showing the process of prioritizing candidate genes for functional screening. To enrich for plausible resistance driver genes, candidates were selected if (1) identified independently by at least two analyses (‘multi-hits’), or (2) implicated in the DDR, or (3) identified as high-impact drivers in the protein-protein network using DriverNet (Bashashati et al., 2012). The latter identifies genes which impact the expression of interacting partners or factors that share the same biological pathway and was used to correct for the fact that frequency analyses integrating data across different omic platforms (‘multi-hits’) fails to identify events restricted to one biochemical domain (e.g., mutation or phosphorylation), but nonetheless important for driving resistance (Bashashati et al., 2012). WE-seq, LCWG-seq and RNA-seq analysis was included for all tumor collections, including tumors whose RAD51-IRIF status was not determined in KB1P(M) (68 for resistant and 43 for naïve tumors), and proteomics and phosphoproteomics data was generated for some of the KB1P(M) tumors (12 for resistant and 12 for naïve tumors).

**(F)** Venn diagram representing the number of genes identified in candidate prioritization analysis. Genes were highlighted if they are previously reported PARPi-resistance factors.

**(G)** Outline of functional genetic enrichment screen. Screen was performed in SUM149PT and RPE1-hTERT *BRCA1^-/-^;TP53^-/-^* cells. Surviving cells were collected after 3 weeks and analysis and hit selection was performed using the MaGECK algorithm (Li et al., 2014)

**(H-I)** Plot of distribution log_2_ratio (fold change (treated versus untreated)) median for all genes versus false discovery rate (FDR) for the screen carried out in SUM149PT (H) and in RPE1-hTERT *BRCA1^-/-^;TP53^-/-^* (I) cells.

**(J)** Kaplan–Meier survival curves of mice transplanted with ORG-KB1P4N.1 tumoroids lines modified with indicated shRNA and treated with 100 mg/kg olaparib. End of treatment (28 days) is indicated by a dotted line.
