## Supplemental Figures for "Multi-omics analysis reveals distinct non-reversion mechanisms of PARPi resistance in BRCA1- versus BRCA2-deficient mammary tumors"

### Supplementary Figure 1 - Related to Figure 1

A

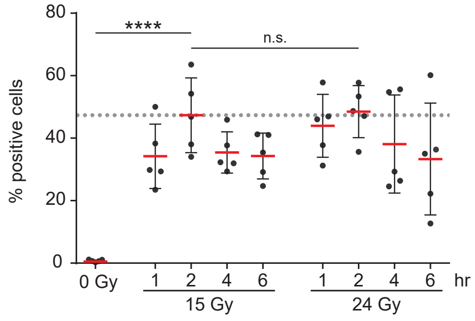


B

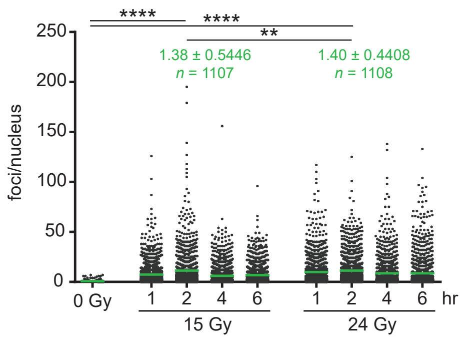


C

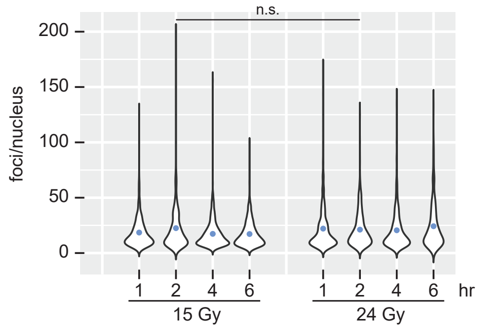


D

| | | mean % positive cells | mean foci/nucleus all cells | mean foci/nucleus positive cells |
|---|---|---|---|---|
| 15 Gy | 1 hr | 34.2 | 7.4 | 18.5 |
| | 2 hr | 47.3 | 11.4 | 22.5 |
| | 4 hr | 35.5 | 6.2 | 17.3 |
| | 6 hr | 34.3 | 6.7 | 17.2 |
| 24 Gy | 1 hr | 44.0 | 9.8 | 22.1 |
| | 2 hr | 48.5 | 11.4 | 21.2 |
| | 4 hr | 38.1 | 8.7 | 20.6 |
| | 6 hr | 33.3 | 8.7 | 24.3 |
| 0 Gy | | 0.51 | 0.1 | n/d |

E

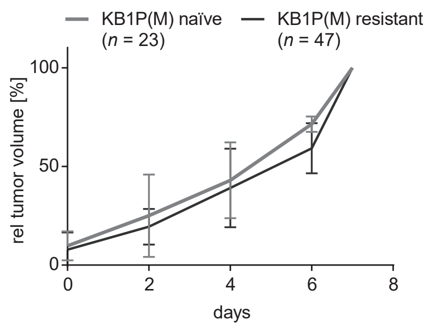


F

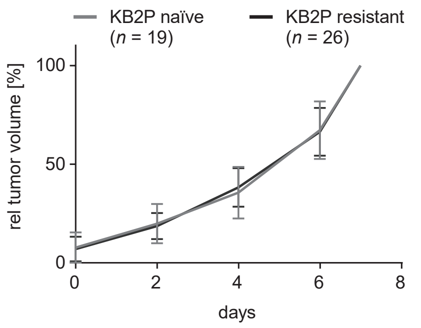


G

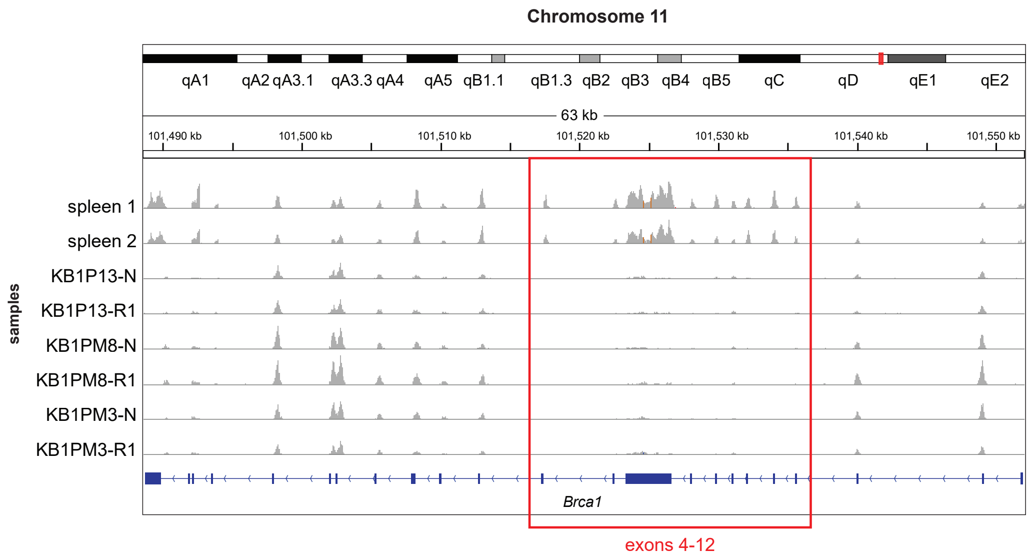


H

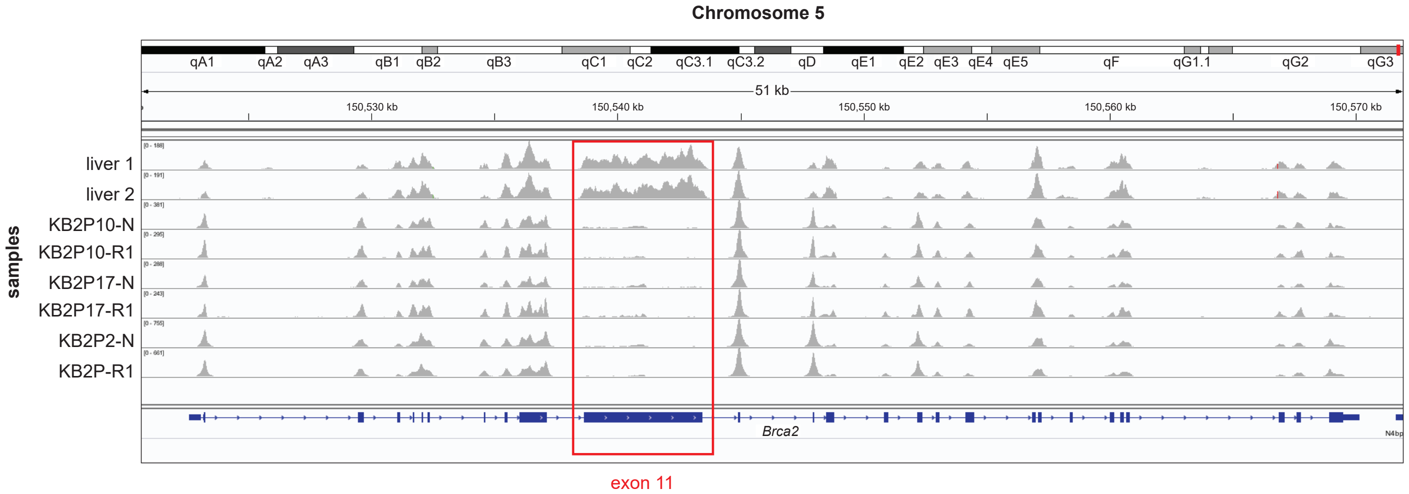

Supplementary Figure 2- Related to Figure 2

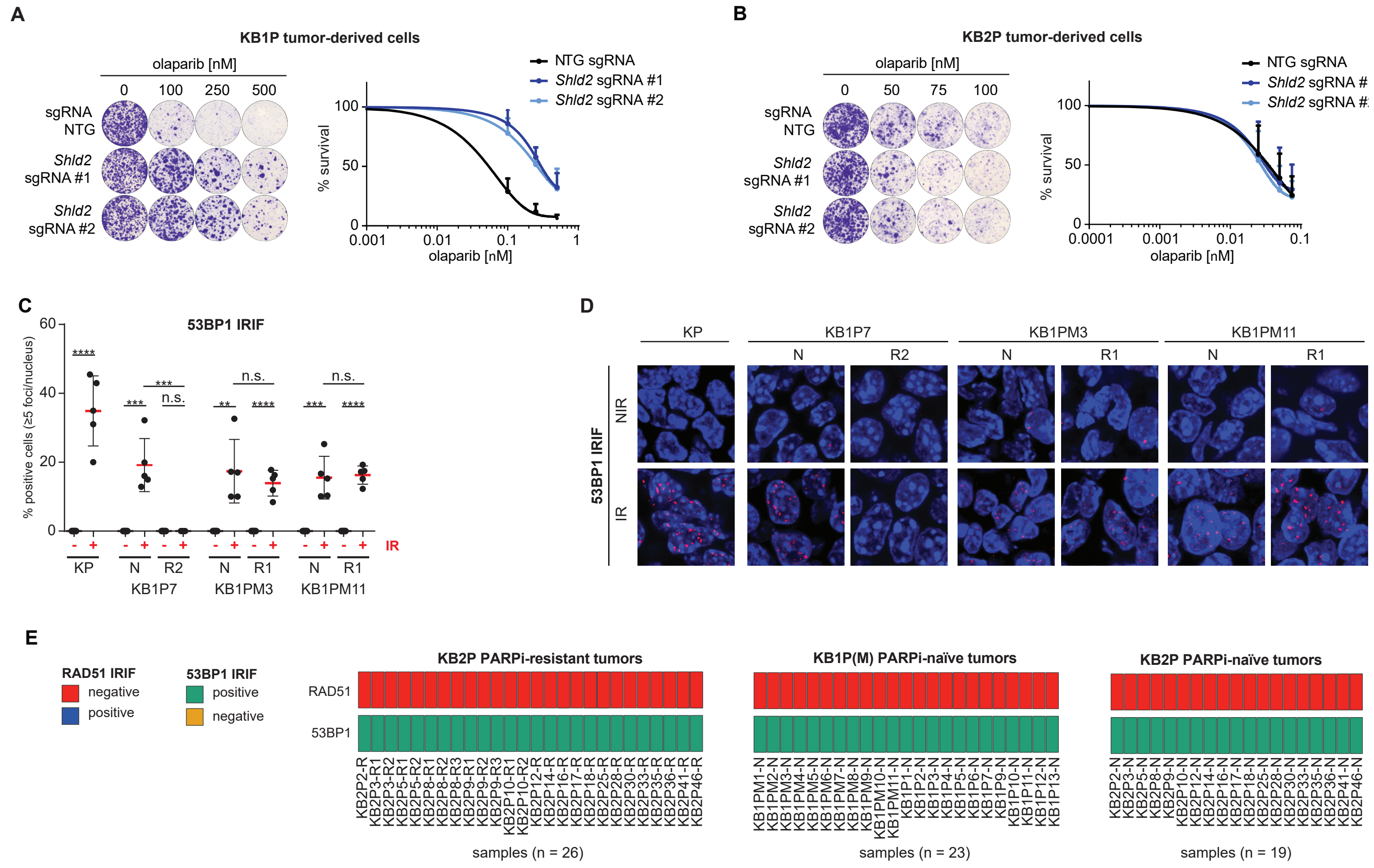

#### Supplementary Figure 3 - Related to Figure 3

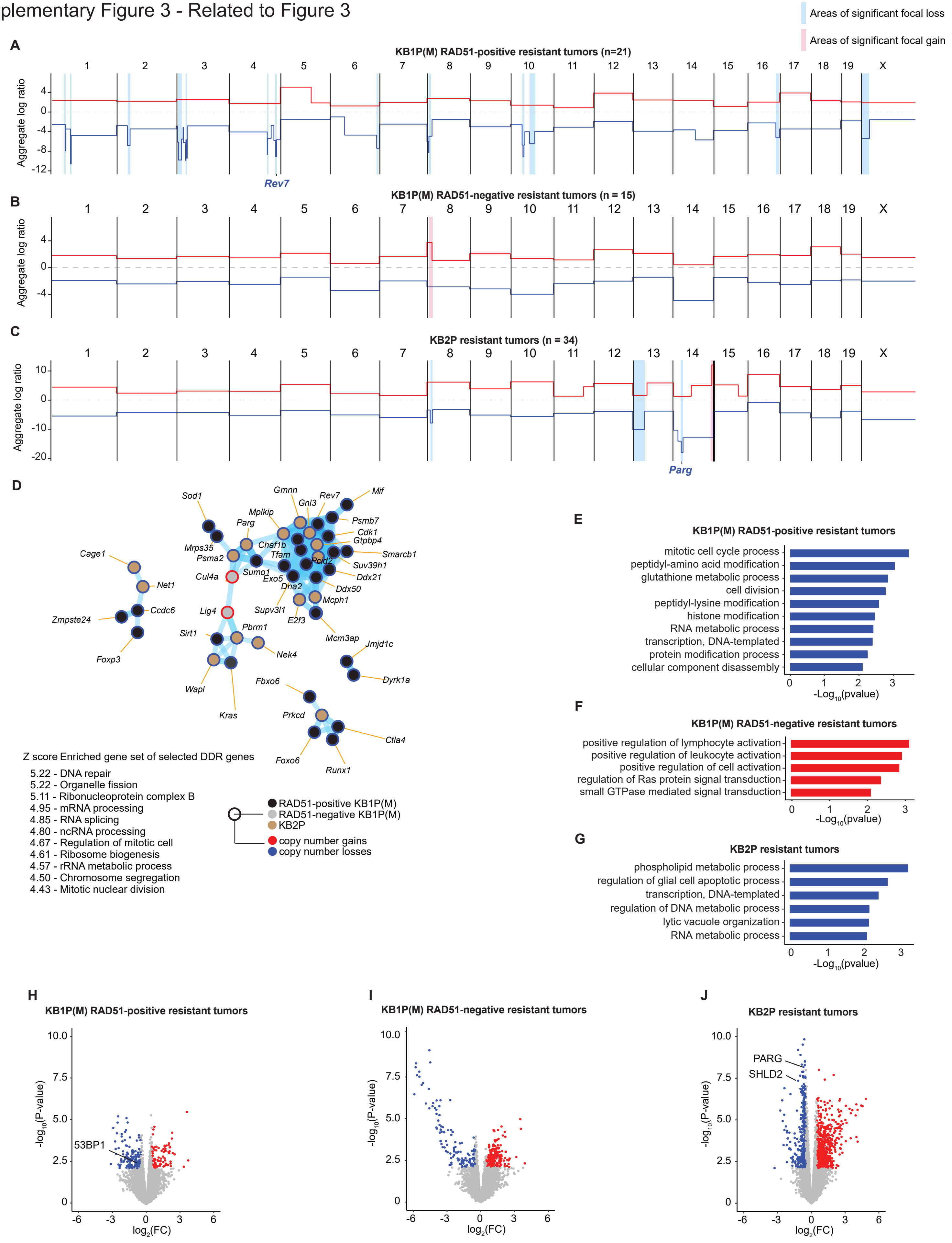

Supplemetary Figure 4 - Related to Figure 4

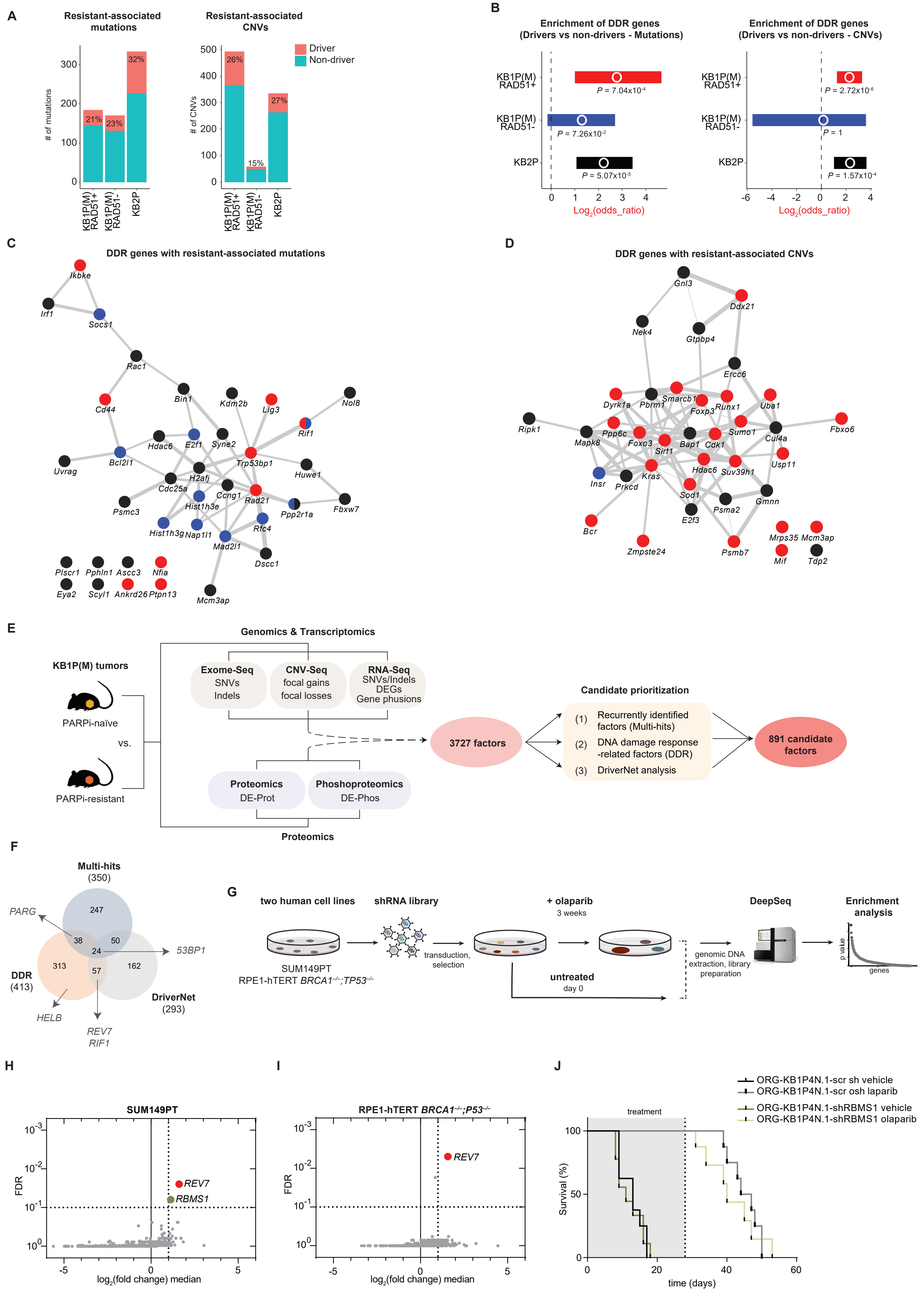
